# Definitive Hematopoietic Stem Cells Minimally Contribute to Embryonic Hematopoiesis

**DOI:** 10.1101/2021.05.02.442359

**Authors:** Bianca A Ulloa, Samima S Habbsa, Kathryn S. Potts, Alana Lewis, Mia McKinstry, Sara G. Payne, Julio Flores, Anastasia Nizhnik, Maria Feliz Norberto, Christian Mosimann, Teresa V Bowman

## Abstract

Hematopoietic stem cells (HSCs) are rare cells that arise in the embryo and sustain adult hematopoiesis. Although the functional potential of nascent HSCs is detectable by transplantation, their native contribution during development is unknown, in part due to the overlapping genesis and marker gene expression with other embryonic blood progenitors. Using single cell transcriptomics, we defined gene signatures that distinguish nascent HSCs from embryonic blood progenitors. Applying a new lineage tracing approach, we selectively tracked HSC output *in situ* and discovered significantly delayed lymphomyeloid contribution. Using a novel inducible HSC injury model, we demonstrated a negligible impact on larval lymphomyelopoiesis following HSC depletion. HSCs are not merely dormant at this developmental stage as they showed robust regeneration after injury. Combined, our findings illuminate that nascent HSCs self-renew but display differentiation latency, while HSC-independent embryonic progenitors sustain developmental hematopoiesis. Understanding the differences among embryonic HSC and progenitor populations will guide improved *de novo* generation and expansion of functional HSCs.

## INTRODUCTION

Hematopoietic stem cells (HSCs) are defined by extensive self-renewal capacity and multilineage differentiation potential. They maintain lifelong hematopoiesis via the production of mature blood cells of the erythroid, myeloid, and lymphoid lineages (Chapple, et al., 2018; Hofer, et al., 2016). The regenerative ability of HSCs to replace a damaged hematopoietic system makes them clinically valuable for hematologic cell replacement therapies (Baron and Storb, 2006). Although more than 50,000 hematopoietic cell transplantations occur worldwide each year (Aljurf, et al., 2019), transplantation is not an option for all patients due to a paucity of appropriate donor cells (Fraint, et al., 2021). Abilities to expand existing donor HSCs or to generate *de novo* cells from less limited stem cell sources such as pluripotent stem cells could conceivably increase transplantation opportunities. Understanding the earliest establishment of HSC self-renewal and multipotency properties could facilitate the development of methods to improve and maximize their therapeutic potential.

HSCs first emerge from the hemogenic endothelium of the newly formed dorsal aorta during embryogenesis (Boisset, et al., 2010; Kissa and Herbomel, 2010). In addition to HSCs, other HSC-independent multi-lineage progenitors emerge in development including erythroid-myeloid progenitors (EMP) and lymphoid-myeloid progenitors (LMP) (He, et al., 2020b; Stachura and Traver, 2016; Chen, et al., 2011). Limited in their self-renewal and differentiation output, these progenitors are mostly regarded as transient in nature (Waas and Maillard, 2017; Stachura and Traver, 2016; Chen, et al., 2011). Seminal work revealed that HSC-independent embryonic progenitors were necessary and sufficient to sustain embryonic hematopoiesis (Chen, et al., 2011). If transient embryonic progenitors can generate erythroid, myeloid, and lymphoid cells, what do HSCs contribute to embryonic/prenatal hematopoiesis? A difficulty in addressing this question is the spatially and temporally overlapping, yet independent generation of HSCs and HSC-independent progenitors and their shared expression of several marker genes used for cell isolation (Zhan, et al., 2018; Tian, et al., 2017; Stachura and Traver, 2016; Chen, et al., 2011; Davidson and Zon, 2004). Consequently, higher resolution data on the distinct transcriptional landscapes of hematopoietic stem and progenitor (HSPC) subsets are needed, in combination with functional assays to examine their endogenous function in development.

Here, we conducted single cell RNA sequencing (scRNA-seq) of newly emerged HSPCs isolated from developing zebrafish and identified seven distinct cell type clusters corresponding to HSCs and three progenitor trajectories. From these studies, we identified temporally dynamic activity of transgenic reporters driven by the regulatory elements of the zebrafish *draculin (drl)* gene that distinguished HSCs from embryonic progenitors. Taking advantage of this difference, we applied a fluorescence-based lineage labeling method to selectively mark HSCs *in vivo* and found that lymphoid and myeloid differentiation from nascent HSCs is significantly delayed as compared to embryonic progenitors. Consistent with this result, we demonstrated a negligible impact on lymphoid and myeloid cell numbers in zebrafish larvae following *in situ* HSC depletion using a novel inducible larval HSC injury model. In contrast, HSCs robustly regenerated following depletion, suggesting they are not completely dormant at this developmental stage. Combined, our data demonstrate that while nascent HSCs possess self-renewal capacity, it is the HSC-independent embryonic progenitors that maintain early developmental hematopoiesis.

## Results

### scRNA-seq revealed distinct embryonic progenitor and HSC clusters

To understand the complexity of HSPC heterogeneity, we used zebrafish as a representative vertebrate model to conduct 10X Genomics scRNA-seq analysis of enriched HSPC populations. We selected cells based on expression of known blood markers *draculin* (*drl*) (Mosimann, et al., 2015; Herbomel, et al., 1999) and *runx1* (Tamplin, et al., 2015).The *drl* gene acts as a pan-lateral plate mesoderm marker from gastrulation to early somitogenesis, and is subsequently one of the earliest markers expressed in the developing hematopoietic system (Mosimann, et al., 2015; Herbomel, et al., 1999). Transgenic *drl* reporters including *drl:mCherry* (driven by the 6.35 kb regulatory region of the *drl* gene) mark both long-lived HSCs and circulating erythrocytes in zebrafish (Prummel, et al., 2019; Henninger, et al., 2017; Robertson, et al., 2016; Mosimann, et al., 2015). To selectively analyze HSPCs, we isolated *drl:mCherry^+^* cells that were negative for the erythroid marker *gata1:GFP* (Traver, et al., 2003; Long, et al., 1997) (Figures 1A, S1A). We analyzed duplicate samples of *drl:mCherry^+^;gata1:GFP^-^* cells isolated from zebrafish embryos at 30 and 52 hours post fertilization (hpf) (referred to hereafter as 1- and 2- days post fertilization (dpf), respectively) to span peak time points of HSC emergence and initial maturation (Henninger, et al., 2017). We also performed scRNA-seq on *runx1+23:nls-mCherry^+^* (referred to hereafter as *runx1:mcherry* or *runx1^+^)* cells isolated from 2 dpf embryos as a secondary marker for validation (Figure S1B). Since expression of *runx1:mCherry* was low at 1 dpf, isolation and scRNA-seq of this population was only possible at 2 dpf.

**Figure 1.**
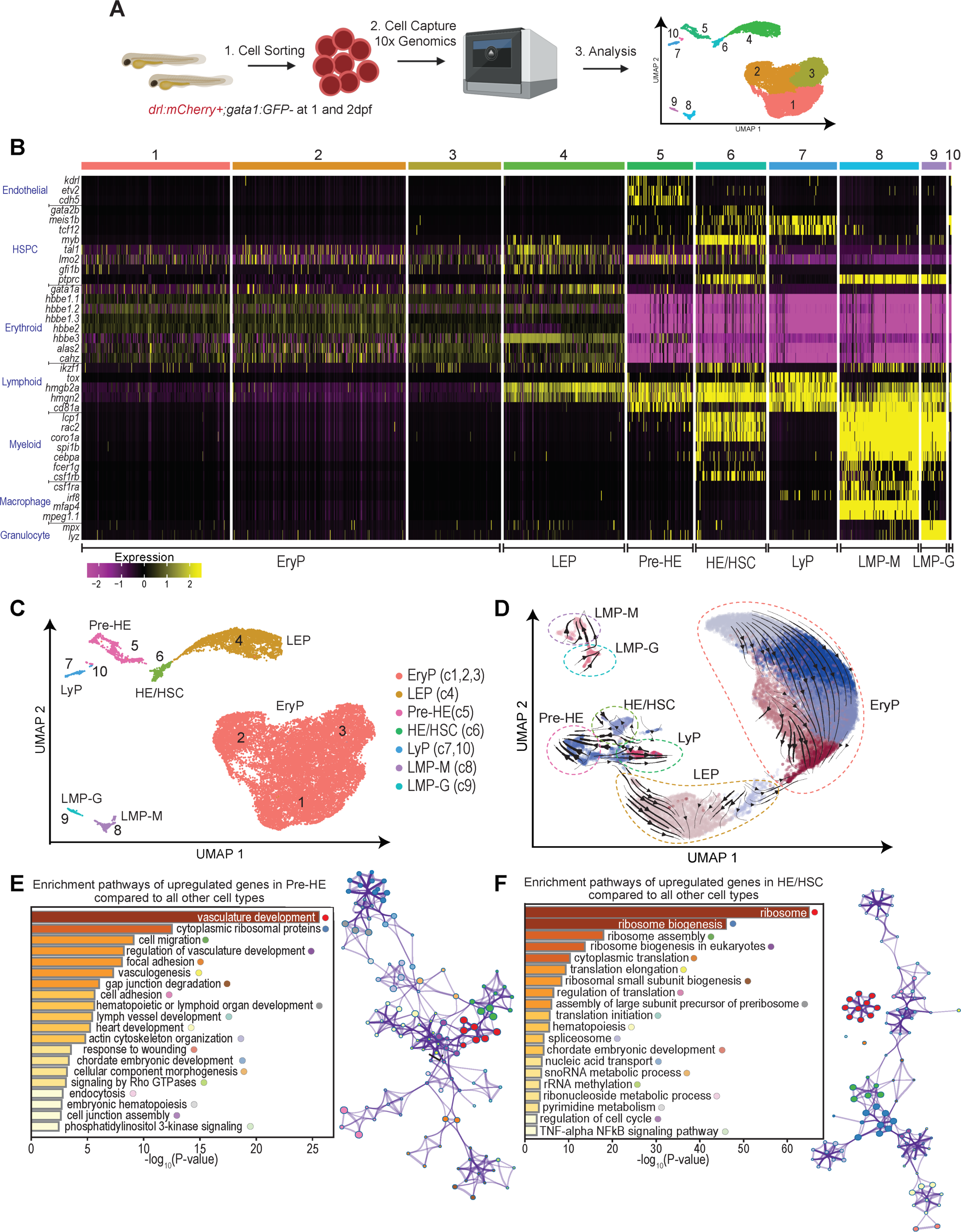
Developmental hematopoietic heterogeneity identified in zebrafish with single cell RNA sequencing. (A) Schema of scRNA-seq experiment: 1) *drl:mCherry*^+^;*gata1:GFP*^-^ cells were isolated by FACS from pools of 1 and 2 dpf embryos to enrich for HSPCs; 2) 10X Genomics single cell RNA libraries were prepared and sequenced; and 3) Downstream computational analysis: UMAP dimensional reduction and unsupervised clustering of *drl:mCherry*^+^;*gata1:GFP*^-^ cells at 1 and 2 dpf with Monocle3 gives ten clusters. (B) Cell type identification of the ten Monocle3 clusters performed using key hematopoietic marker genes supported by previous zebrafish, murine, and human literature. Cell types identified: erythroid progenitors (EryP), Clusters 1-3 (c1,2,3); lymphoerythroid progenitor (LEP), Cluster 4 (c4); pre-hemogenic endothelium (Pre-HE), Cluster 5 (c5); hemogenic endothelium giving rise to hematopoietic stem cells (HE/HSC), Cluster 6 (c6); lymphoid progenitor (LyP), Cluster 7 and 10 (c7,10); lymphomyeloid progenitor with higher macrophage marker expression (LMP-M), Cluster 8 (c8); and lymphomyeloid progenitor with higher granulocyte marker expression (LMP-G), Cluster 9 (c9). Expression bar: scaling is performed per gene where mean is close to 0 and standard deviation of 1. Only genes with the highest scaled expression value will show the brightest yellow color in any of the cells (standard deviation above 2), and those that have the lowest scaled expression will be purple (standard deviation below -2). (C) The cell type classifications in UMAP space as identified in Fig. 1B. (D)RNA velocities overlaid over UMAP clusters of *drl:mCherry^+^gata1:GFP^-^* cells at 1 and 2 dpf. The different colors represent the Seurat clusters derived from reprocessing the data. The dotted lines encircling each cluster are cell types identified with previously used marker genes (refer to Fig S1D). Arrows indicate inferred differentiation trajectory as determined by RNA velocities. (E-F) Enrichment pathway analysis of upregulated top marker genes in Pre-HE (E) and HE/HSC (F) clusters, conducted using Metascape. Top 200 markers selected with the following criteria: fraction expressing >10% and marker test p value < 0.00005. The colored circles after each term represent a colored dot on the network representation to the right of the significance bar plot. See also Figure S1.

Summary statistics of quality control for analyzed cell populations can be found in Supplemental Table 1. Subsequent analysis was performed using established tools, Seurat and Monocle3. After batch-correction by matching mutual nearest neighbors on Monocle3 (Haghverdi, et al., 2018), we used uniform manifold approximation and projection (UMAP) to reduce the dimensionality of our data (Hao, et al., 2020; Cao, et al., 2019; McInnes, 2018). Focusing on 1 and 2 dpf *drl^+^gata1^-^* cell populations, we conducted unsupervised clustering and identified 10 major clusters, in line with a heterogeneous HSPC pool at these developmental stages (Figure 1A). Comparison of the 2 dpf *drl^+^gata1^-^* and *runx1*^+^ scRNA-seq data revealed a significant overlap between the populations identified, confirming that our observed heterogeneity within the embryonic HSPC pool was not a consequence of the markers used, but rather an indication of the complexity of developmental hematopoiesis (Figure S1C).

We designated the identities of the cell clusters based on signature marker gene expression as defined in zebrafish and murine developmental hematopoiesis (Figures 1B-C). C5 was designated as pre-hemogenic endothelial cells (Pre-HE) as it featured strong expression of pan-endothelial genes (*kdrl, etv2, cdh5*) (Bonkhofer, et al., 2019; Oh, et al., 2015; Sauteur, et al., 2014), minimal expression of HE marker genes (*lmo2, tal1, meis1b*) (Wang, et al., 2018; Patterson, et al., 2007), and absent expression of HSC genes *gata2b* (Butko, et al., 2015) and *myb* (Zhang, et al., 2011). Thus, C5 represents a cell population initiating an endothelial-to-hematopoietic transition (Zovein, et al., 2008). In contrast, C6 cells had high expression of key hemogenic and HSC marker genes including *gata2b, meis1b*, and *myb*, as well as expression of lineage-restricted genes indicative of multipotent priming such as *gata1a* (erythroid) (Gutiérrez, et al., 2020; Galloway, et al., 2005), *ikzf1* (lymphoid) (Huang, et al., 2019; Willett, et al., 2001), and *coro1a* and *csf1rb* (myeloid) (Li, et al., 2012; Carstanjen, et al., 2005; Song, et al., 2004); we consequently designated cluster C6 as hemogenic endothelial cells generating hematopoietic stem cells (HE/HSC). The C4 cells expressed erythroid and lymphoid markers with low expression of myeloid markers, suggesting this population is analogous to the recently described lympho-erythroid-primed progenitor (LEP) in zebrafish (Kasper, et al., 2020). Cells with lympho-myeloid progenitor (LMP) expression profile occurred in two clusters: an LMP population with higher macrophage marker expression (*csf1ra, irf8, mfap4, mpeg1.1*) (Rojo, et al., 2019; Li, et al., 2011; Zakrzewska, et al., 2010) (LMP-M, C8) and an LMP population with higher granulocyte marker expression (*mpx, lyz*) (Hall, et al., 2007; Renshaw, et al., 2006) (LMP-G, C9). The C7 and C10 clusters were highly similar and expressed progenitor-associated genes including *meis1b* and *tcf12*, and mainly lymphoid-associated genes including *tox* and *cd81a*, suggesting C7 and C10 represent more restricted lymphoid progenitors (LyP). Most cells resided in clusters C1, C2, and C3 and predominantly expressed erythroid markers, which we broadly designated as erythroid progenitors (EryP). Together, our identified clusters encompass the range of HSPC-associated progenitor traits (Hou, et al., 2020; Kasper, et al., 2020; Zhu, et al., 2020; Xue, et al., 2019; Zhou, et al., 2016), here resolved into distinct progenitor clusters.

We inferred the developmental lineage trajectories of the different cell types from calculated RNA velocities using scVelo (Bergen, et al., 2020) (Figure 1D). We identified the same cell type classifications using this analysis as we observed with the original Monocle3 clustering (Figure S1D). RNA velocity projections documented stochastic cell states between the LMP-M and LMP-G cells with a separation between LMPs and the other populations (Figure 1D). In this cluster projection, the Pre-HE is a common precursor giving rise to three potential differentiation trajectories: HE/HSC, LyP, or LEP, suggesting a developmental connection among these cell states. The data also support a differentiation trajectory of a subset of LEPs to become more restricted EryPs. These progressions are further supported by pseudotime analysis using Monocle3 (Cao, et al., 2019; Trapnell, et al., 2014) (Figure S1E). To give directionality to the pseudotime trajectory, we defined the Pre-HE as the classical starting point of differentiation. Based on this analysis, the LEP and EryP represent more committed differentiated states.

To gain insights into functional features of the transcriptomes defining our ten clusters, we took the most highly and selectively expressed marker genes (fraction of cells expressing the gene within the cell group >10% and marker test p value < 0.00005) for each cluster and conducted gene network enrichment analysis using Metascape and Ingenuity Pathway Analysis (IPA) (Figures 1E-F, Supplemental Table 2) (Zhou, et al., 2019). The Pre-HE cluster genes were enriched for terms related to vasculature development, cytoplasmic ribosomal proteins, cell migration, and hematopoiesis (Figure 1E). HE/HSC cluster genes showed enrichment in ribosomal biogenesis and tumor necrosis factor alpha signaling via nuclear factor κB signaling pathway, which has been associated with HSC development (Espin-Palazon, et al., 2014) (Figure 1F). The LyP cluster genes showed strong enrichment for dorsal aorta development, Notch signaling pathway, lymphoid organ development, and regulation of cell differentiation (Supplemental Table 2). As predicted, the LMP populations were enriched for both myeloid and lymphoid pathways, such as for myeloid leukocyte differentiation/activation, macrophage chemotaxis, and T-cell receptor signaling pathway. Lastly, LEP and EryP populations were both enriched for erythrocyte signatures as well as cell cycle. Taken together, gene set enrichment network analysis supports our cell type classifications and highlight important signatures for HSPC development and lineages as captured in our scRNA-seq dataset.

HSPC development is highly conserved between zebrafish and mammals (Avagyan and Zon, 2016; Robertson, et al., 2016). To assess the similarity of the HE/HSC gene signatures in zebrafish and mammals, we compared our data to gene signatures defined in recently published HE and pre-HSC/HSC scRNA-seq experiments from the analogous developmental time points in mice (Hou, et al., 2020; Vink, et al., 2020; Zhu, et al., 2020; Zhou, et al., 2016) (Supplemental Table 3). We evaluated the relative aggregate gene expressions (gene-set score) of murine pre-HE (Zhu, et al., 2020) and HE/pre-HSC/HSC signatures (Hou, et al., 2020; Vink, et al., 2020; Zhu, et al., 2020; Zhou, et al., 2016) across our zebrafish HSPC clustering (Figures S1F-G). The murine pre-HE gene-set score faithfully recapitulated our classification of the zebrafish pre-HE cluster (Figure S1F). Although the murine HE/HSC gene-set score had generalized coverage throughout the zebrafish cell types the highest values were in the HE/HSC zebrafish cluster (Figure S1G). The HE/HSC gene-set score indicates that many of the genes expressed in this population are also expressed by other cell types and suggests our progenitor populations (LEPs, LMPs, EryPs, and LyPs) arose from HE cells. The LMP population (LMP-G and LMP-M) also had a relatively high gene-set score for both pre-HE and HE murine signatures, which could again suggest their pre-HE/HE origin. Overall, our results support that our zebrafish pre-HE and HE/HSC clusters are conserved with previous mammalian scRNAseq signatures and illustrates the complexity of deciphering the heterogeneous pre-HSC/HSC compartment at this early developmental window.

### Temporal regulation of the *drl:mCherry* reporter distinguishes HSPC subsets

We posited that the overlapping waves of HSPCs (Tian, et al., 2017; Stachura and Traver, 2016; McGrath, et al., 2015) may lead to differences in their prevalence over time. Therefore, we investigated the HSPC composition over the 1 and 2 dpf time frame (Figure 2A). We collected a greater number of EryP and HE/HSCs at 2 dpf as compared to 1 dpf (Figure 2B, Supplemental Table 4). The EryP population also far outnumbered any of the other cell types at 2 dpf; in contrast, we observed a substantial decrease in the number of Pre-HE cells, LyPs, LEPs and LMPs at this later time point. In the *drl:mCherry* fish, expression of *mCherry* is a surrogate estimate for the *drl* regulatory element activity. Upon examination of the *drl:mCherry* transgene expression, we noticed high *mCherry* transcript levels in several HSPC subsets at 1 dpf, including Pre-HE, HE/HSC, EryP, and LEP, but this became largely restricted to the HE/HSC subset by 2 dpf. Based on these data, we posited that we could exploit the temporal differences in *drl*-driven transgene expression as a tool to distinguish HSCs from other progenitors.

**Figure 2.**
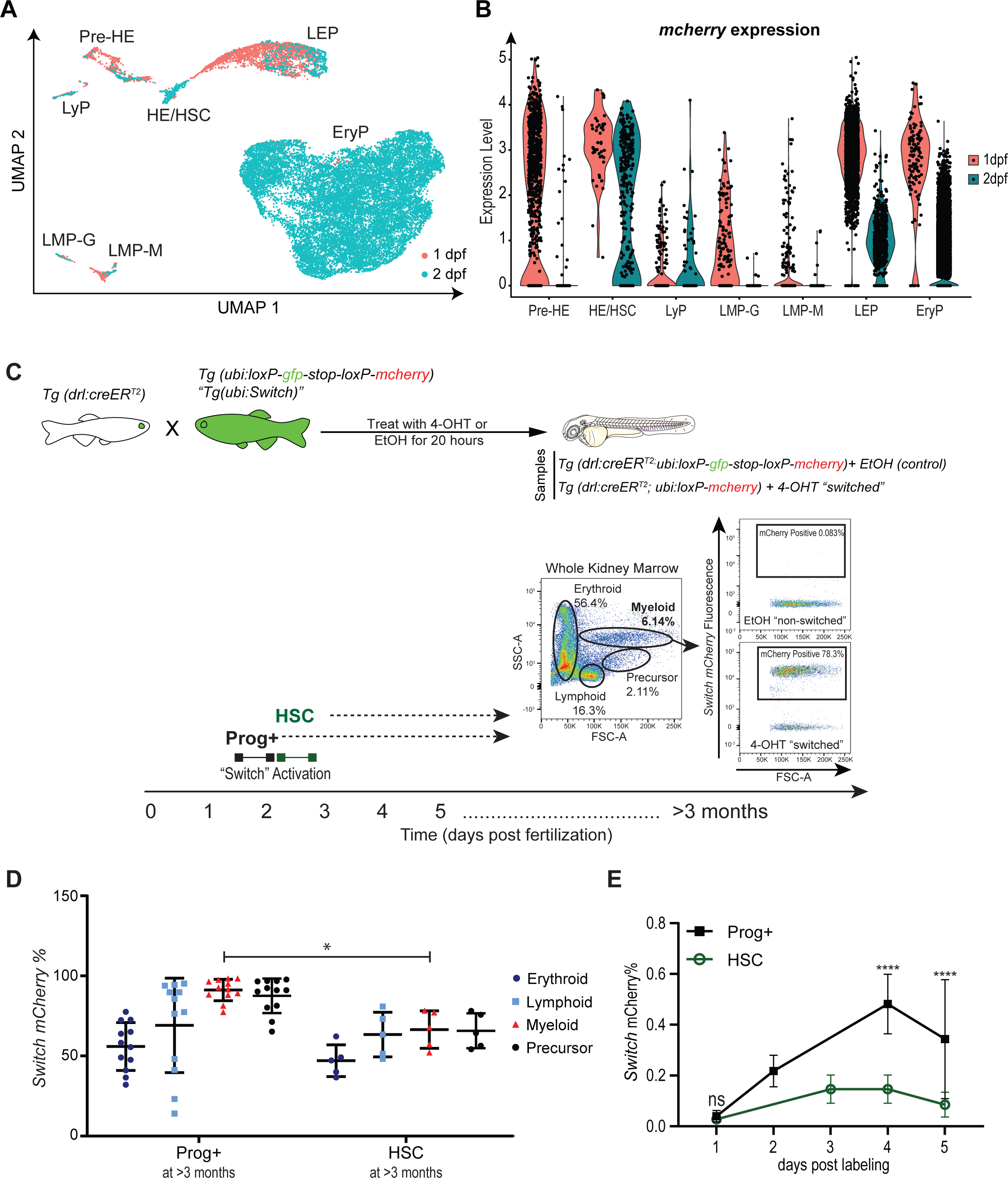
Dynamic regulation of the *drl* promoter distinguishes HSPC subsets. (A) UMAP dimensional reduction and unsupervised clustering of *drl:mCherry*^+^;*gata1:GFP*^-^ cells at 1 and 2 dpf showing their original collection time point. (B) *mCherry* expression level driven by the *drl* promoter for the different cell types collected at 1 and 2 dpf. (C) Schematic illustrating *drl*-specific, 4-OHT-inducible lineage tracing system. *Tg(drl:creER^T2^)* homozygous transgenic fish were crossed with hemizygous *ubi:loxP-gfp-stop-loxP-mCherry* (*ubi:Switch*). Zebrafish embryos were exposed to 12 µM 4-OH-Tamoxifen (4-OHT) or ethanol (EtOH) vehicle control for 20 hours starting at either 30 hpf (labeled as Prog+, which includes embryonic progenitors and HSCs) or 54 hpf (labeled as HSC). Those embryos exposed to 4-OHT induce Cre recombination of *loxP* sites, leading to a permanent “switch” excising the *GFP* cassette and resulting in expression of mCherry fluorescence. The resulting mCherry^+^ cells are the descendants of Prog+- and HSC-labeled cells. Representative flow cytometry analysis: forward (FSC) and side scatter (SSC) parameters were used to define the major blood cell populations (erythroid, myeloid, lymphoid, and precursor) in whole kidney marrow (WKM) (>3 months). The frequency of mCherry^+^ cells within each cell lineage was then calculated. Example flow cytometry plots (right) of mCherry contribution to the myeloid population. Gates were set based on ethanol treated (EtOH) “non-switched” controls. (D) Quantification of mCherry^+^ percentage from experimental groups Prog+ and HSC within each blood cell lineage found in WKM at >3 months post fertilization (zebrafish adulthood). Data points represent individual zebrafish WKM with mean ± standard deviation (N = 5-12). (E) Quantification of mCherry^+^ percentage at 1-5 days post labeling in Prog+ and HSC labeled cohorts. N = 5-11 samples, 7-10 pooled larvae per sample. Two-way ANOVA with Sidak’s multiple comparison test was used for analysis (mean± standard deviation). *p-value < 0.05, ****p-value *≤* 0.0001. See also Figure S2.

The *drl* regulatory elements consist of an early pan-lateral plate mesodermal enhancer and two later-acting, cardiovascular-specific regulatory elements that confine *drl* reporter expression to the heart, endothelium, and blood lineages starting in late somitogenesis (Prummel, et al., 2019). Previously, the *Tg(drl:creER^T2^)* transgenic zebrafish that permits 4-OH-Tamoxifen (4-OHT)-inducible Cre activation was used to demonstrate that, in hematopoiesis, embryonic *drl*-expressing cells contribute to lifelong blood production (Henninger, et al., 2017). Only adult hematopoiesis was examined in that study, leaving the differentiation dynamics and lineage contributions of *drl-*expressing hematopoietic lineages during development untested. Taking advantage of this system, we performed a fluorescence-based lineage-tracing method to distinctly label progenitors and HSCs at 1 and 2 dpf and track their lineage output. In double transgenic *Tg*(*drl:creER^T2^*;*ubi:loxP-GFP-loxP-mCherry*) (referred to hereafter as *drl:creER^T2^;ubi:Switch*) zebrafish, 4-OHT-inducible Cre recombination removes the *GFP* cassette from *ubi:Switch* in *drl* reporter-expressing cells, leading to permanent *mCherry* expression in *drl^+^* labeled cells and their progeny (Mosimann, et al., 2015; Mosimann, et al., 2011) (Figure 2C). We then monitored mCherry-fluorescent cells via flow cytometry and fluorescence imaging to determine the contributions of distinct *drl^+^* HSPCs to developmental and adult hematopoiesis.

To induce recombination, we exposed embryos to either 4-OHT or ethanol (EtOH) vehicle control starting at 30 hpf to label embryonic hematopoietic progenitors and HSCs (hereafter referred to as Prog+), and at 54 hpf to label HSCs as per our scRNA-seq analysis (Figures 2B-C). Consistent with previous findings (Henninger, et al., 2017), we validated that our lineage tracing system was labeling long-term multipotent HSCs in both Prog+ and HSC cohorts by confirming the presence of mCherry^+^ progeny in erythroid, lymphoid, and myeloid cells in 3-4 month old adult kidney marrow cells via flow cytometry (Traver, et al., 2003) (Figure 2C-D). We compared the recombination efficiency one day post- 4-OHT exposure and demonstrated a similar mCherry^+^ percentage of Prog+- and HSC-labeling, indicating similar Cre recombination efficiency at both switch time points (30 and 54 hpf) that allows direct comparison of the output of Prog+ and HSC lineage tracing in embryos (Figure 2E). By comparing Prog+ and HSC mCherry^+^ cell frequencies using flow cytometric and fluorescence imaging quantification, we discovered significant differences in the expansion of mCherry^+^ progeny from Prog+ *versus* HSCs over time, such as Prog+ progeny significantly outnumbering HSC progeny at 4 and 5 dpf (Figure 2E, S2). This disparity suggests a potential difference in the contribution of HSC *versus* Prog+ progeny to developmental hematopoiesis that we more closely inspected in the next experiments.

### Larval lymphoid and myeloid differentiation kinetics are distinct between HSCs and other embryonic progenitors

Based on the classical hematopoietic hierarchy (Stachura and Traver, 2016), we expected that mCherry^+^ progeny from HSC and Prog+ lineage tracing could either be replicative copies of the original labeled cell and/or differentiated mature lymphoid, myeloid, or erythroid cells. We focused on lymphoid and myeloid lineages and excluded analysis of erythroid cells because the *drl* promoter also directly labels mature erythrocytes, thus making it unclear if the lineage tracing in erythrocytes was HSC/Prog+ or erythrocyte derived (Henninger, et al., 2017; Robertson, et al., 2016; Mosimann, et al., 2015). During vertebrate development, distinct hematopoietic cell types reside in particular anatomical locations at specific time points (Stachura and Traver, 2016). For example, erythrocytes and thrombocytes travel rapidly in circulation (Brönnimann, et al., 2018; Khandekar, et al., 2012), macrophages and dendritic cells can reside in the skin (Zhan, et al., 2018), neutrophils ubiquitously spread throughout circulation, epidermis, or congregating in stressed tissue (Le Guyader, et al., 2008), and T-cells are abundant in the thymus (Tian, et al., 2017). To measure Prog+ and HSC contribution to T-lymphocytes, we imaged mCherry fluorescence in the thymus in 5-16 dpf larvae (Figure 3A). Prog+ cells contributed to thymic cells by 5 dpf (Figures 3B, 3D, S3). In contrast, HSC contribution to the thymus was significantly delayed until after 7 dpf (Figures 3C-D, S3). If the difference in lymphoid contribution was simply due to the one-day delay in Cre activation, then we would expect a one-day delay in contribution between Prog+ and HSCs. Instead, we observed more than a two-day delay with thymus contribution not reaching a comparable level until 10 dpf, suggesting a functional difference in lymphoid differentiation between Prog+ and HSCs.

**Figure 3.**
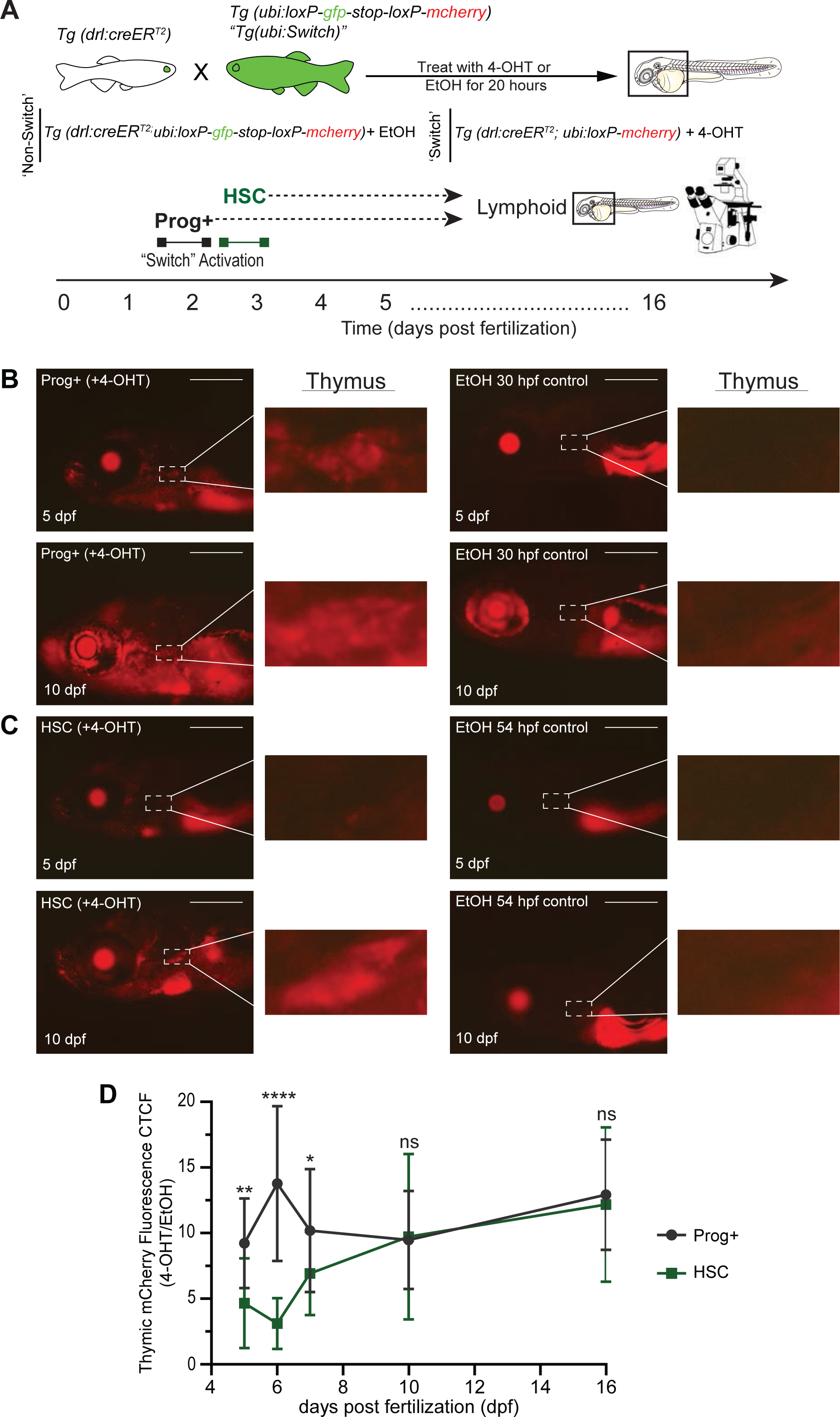
Thymic T-cell contribution by embryonic progenitors and HSCs is distinct. (A) Experimental schema of 4-OHT-inducible lineage tracing to examine larval T-cell production in the thymus from Prog+- and HSC-labeled cohorts. (B-C) Fluorescent images of 4-OHT-induced switch (left) and ethanol non-switched controls (right) *Tg(drl:creER^T2^; ubi:Switch)* larvae for (B) Prog+ and (C) HSC populations. Top images are 5 dpf larvae and bottom images are 10 dpf larvae. Dashed box and inset showing the thymus where T-cells colonize. mCherry^+^ fluorescence corresponds to *drl+* switched daughter cells. Scale bar = 500 µm. Representative images shown, quantification in D. (D) Quantification of mCherry fluorescence intensity in the thymic region in larvae of Prog+- and HSC- labeled cohorts measured over a time course of 5-16 dpf. Mean ± standard deviation of the mCherry^+^ corrected total cell fluorescence (CTCF) at each time point is shown. Two-way ANOVA with Sidak’s multiple comparison (N = 6-30 per larvae/day). *p-value *≤* 0.05, **p-value *≤* 0.01, ****p-value *≤* 0.0001. See also Figure S3.

To examine HSC and Prog+ contribution to myeloid cells, we combined our lineage labeling system with a *Tg(mpx:GFP)* (Renshaw, et al., 2006) line to mark granulocytes, generating triple-transgenic *drl:CreER^T2^;mpx:GFP;ubi:Switch)* zebrafish (Figure 4A). Fluorescence imaging of *Tg(mpx:GFP;ubi:Switch)* demonstrated that *mpx:GFP^+^* cells were brighter than the more diffuse *ubi:Switch* GFP signal (Figure 4B). Flow cytometry analysis confirmed that *drl:creER^T2^;mpx:GFP;ubi:Switch* larvae contain a unique GFP^high^ fraction that is absent in *drl:CreER^T2^;ubi:Switch* but with the same fluorescence intensity as *mpx:GFP*^high^ cells in *mpx:GFP* single transgenics (Figures 4C, S4). Thus, the mCherry^+^ cells within this *mpx:GFP*^high^ fraction represent myeloid progeny from *drl+* cells (Figure 4C). Using this system, we found that HSC contribution to granulocytes was not appreciable until after 7 dpf, while Prog+ cells robustly generated myeloid progeny by 5 dpf (Figure 4D). As seen in our lymphoid results, myeloid contribution by Prog+ and HSCs reached comparable levels by 10 dpf. Consistent with a distinct function of Prog+ cells and HSCs, the lag in generating granulocytes was again greater than the expected one-day difference if simply a consequence of delayed Cre activation. Together, these data demonstrate that embryonic and early larval hematopoiesis, including T-cells and granulocytes, are sustained by an embryonic progenitor pool and not by HSCs.

**Figure 4.**
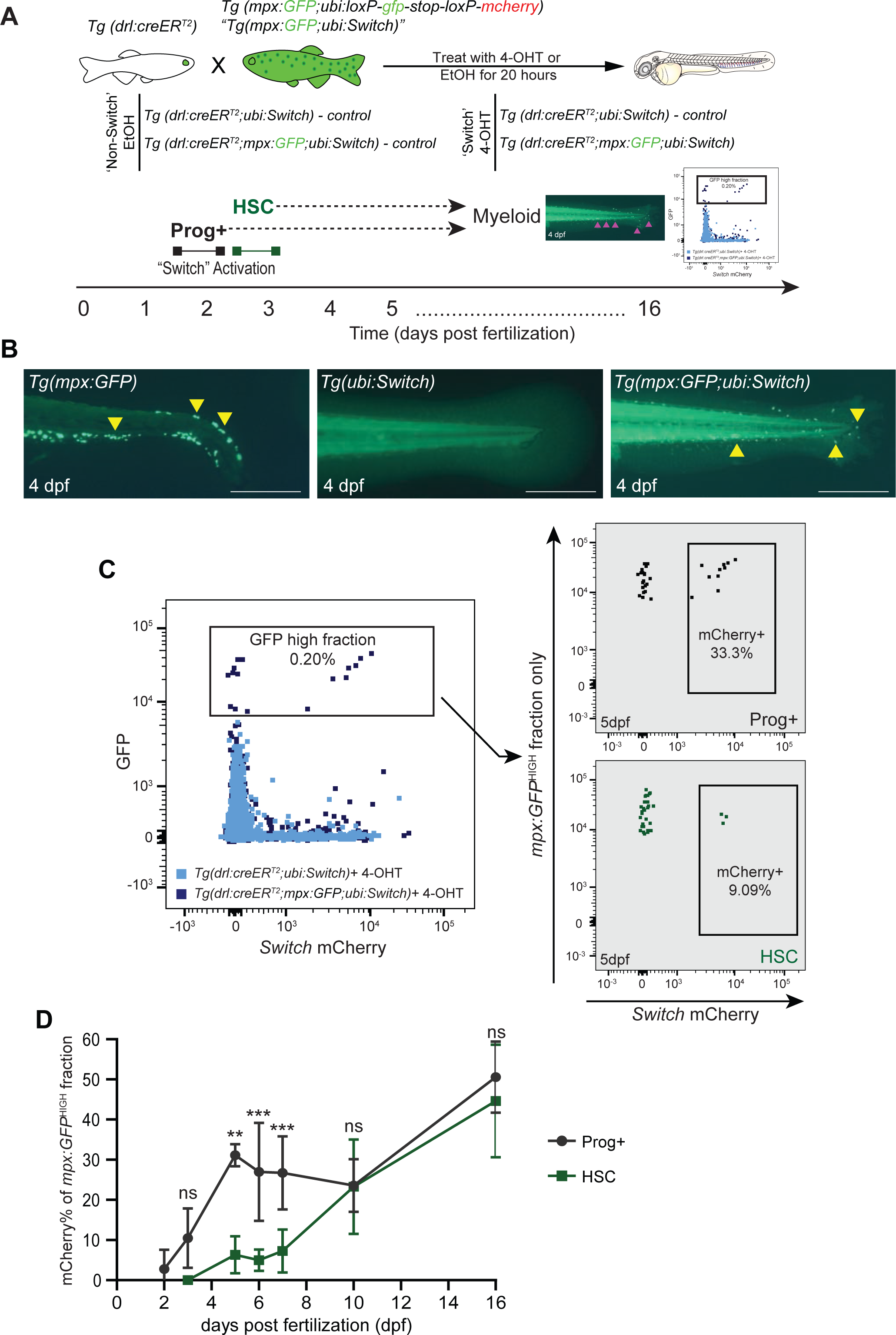
Granulocyte contribution by embryonic progenitors and HSCs is distinct. (A) Experimental schema of 4-OHT-inducible lineage tracing to examine larval granulocyte production from Prog+- and HSC-labeled cohorts. Granulocytes are distinguished by high GFP expression from the *mpx:GFP* transgene compared to the low GFP intensity of *ubi:Switch*. (B) Fluorescent images of the tail region of 4 dpf *Tg(mpx:GFP)* (left), *Tg(ubi:Switch)* (middle), and *Tg(mpx:GFP;ubi:Switch)* (right). *mpx:GFP^+^* granulocytes are denoted by yellow arrowheads. Scale bar = 500 µm. (C) Overlay flow cytometry plot (left) showing higher levels of *mpx:GFP* signal in *Tg(drl:creER^T2^;mpx:GFP;ubi:Switch)* (dark purple) compared to *Tg(drl:creER^T2^;ubi:Switch)* control (light blue). The frequency of mCherry^+^ switch cells was then calculated in the GFP^high^ fraction in Prog+- and HSC-labeled cohorts. Representative flow cytometry plots from 5 dpf larvae (right). (D) Quantification of percentage of mCherry^+^ cells within the *mpx:GFP^high^* fraction in Prog+ and HSC populations measured over a time course of 2-16 dpf (N = 3-7 samples, 7-10 fish per sample). Mean ± standard deviation of the mCherry^+^ percentage at each time point is shown. Two-way ANOVA with Sidak’s multiple comparison was used for this analysis. **p-value *≤* 0.01, ***p-value *≤* 0.001. See also Figure S4.

### HSCs regenerate following induced larval hematopoietic injury

Our lineage-tracing data suggest that HSCs do not significantly contribute to larval lymphomyelopoiesis (Figures 3-4). To assess if nascent HSCs are completely dormant at these early developmental stages, we developed a novel *in situ* larval hematopoietic regeneration assay. Adult HSC regenerative properties are routinely measured following myeloablative injuries such as chemotherapy or irradiation. Such approaches are not employable during development as all tissues are highly proliferative and thus susceptible to cell death following these treatments. The nitroreductase/metronidazole (NTR/MTZ) system is commonly used in zebrafish to direct conditional and inducible cell ablation (Curado, et al., 2008). NTR is a bacterial enzyme that metabolizes the pro-drug MTZ into a toxic DNA cross-linking intermediate leading to apoptotic cell death. We generated transgenic zebrafish that express a Cyan fluorescent protein (CFP)-NTR fusion under the control of the *drl* regulatory elements to drive observable NTR expression during early developmental hematopoiesis and to enable temporally-controlled HSC depletion (Figure 5A) . Harnessing the positional variability in Tol2-based transgene activity common to zebrafish transgenesis (Suster, et al., 2009), we selected the resulting *Tg(drl:CFP-NTR)* (subsequently called *drl:CFP-NTR*) for high hematopoietic expression and comparatively low activity in other *drl*-expressing lineages such as the heart to minimize MTZ-induced cell ablation outside hematopoietic lineages. We confirmed the overlap between *drl:mCherry* and *drl:CFP-NTR,* demonstrating faithful expression of the *CFP-NTR* transgene (Figure S5A).

**Figure 5.**
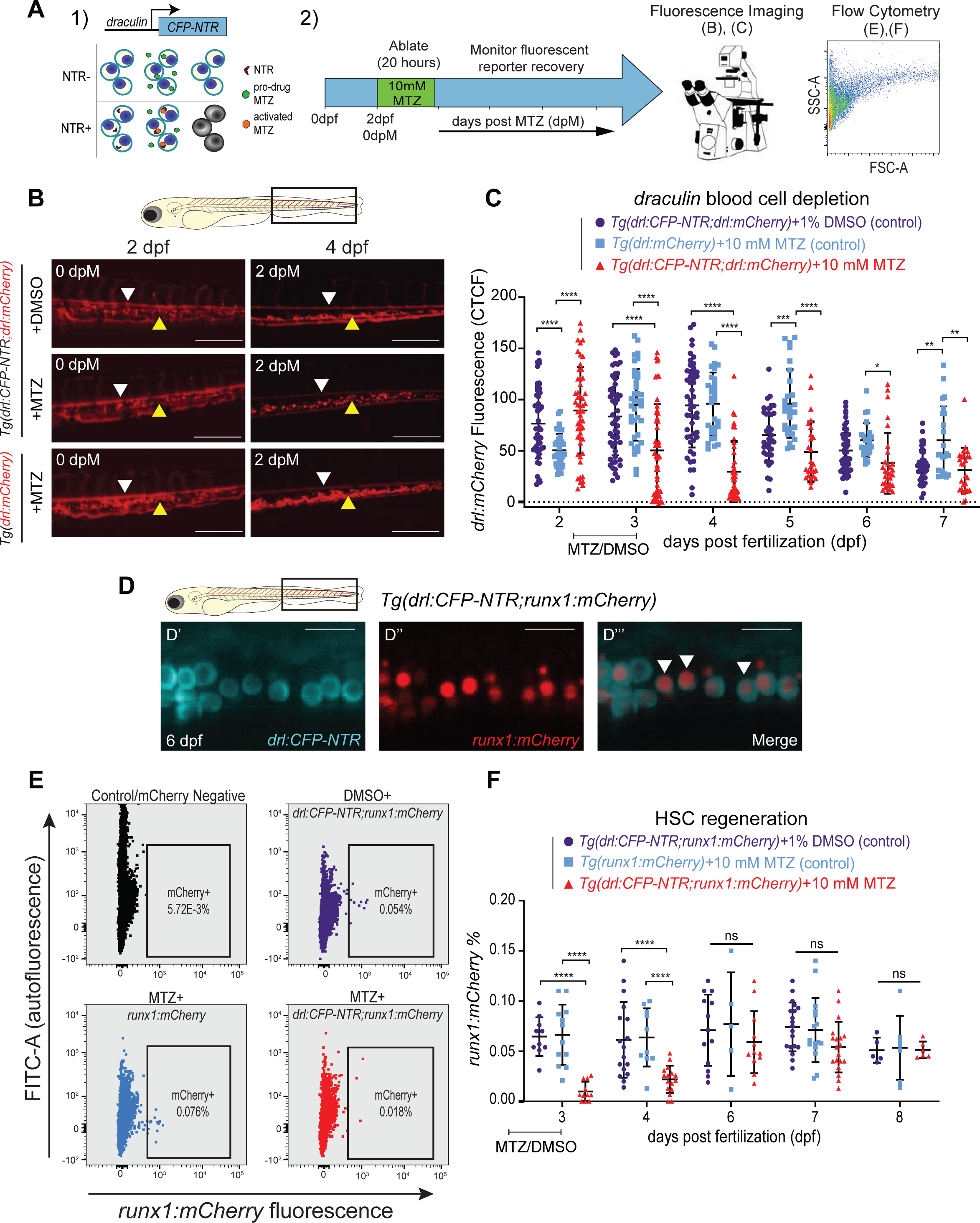
HSCs regenerate following their targeted depletion in early development. (A) Experimental schema of regeneration assay: 1) *Mechanism*: *drl* promoter drives expression of a *CFP-NTR* (Nitroreductase) transgene. NTR converts metronidazole (MTZ) into a toxic intermediate that triggers apoptosis of only *drl:CFP-NTR*-expressing cells. 2) *Timeline*: *Tg(drl:CFP-NTR)* or control larvae were treated with 10 mM MTZ or 1% DMSO vehicle control for 20 hours from 54-74 hours post-fertilization (∼2-3 dpf) to specifically target HSCs, and were then monitored for their recovery using fluorescence imaging and flow cytometry. (B) Fluorescent images of *Tg(drl:CFP-NTR^+^;drl:mCherry^+^)* and *Tg(drl:mCherry^+^)* embryos treated with 10 mM MTZ or 1% DMSO (control) at 2 and 4 dpf (0 and 2 days post MTZ [dpM], respectively). Arrowheads indicate remaining stationary (yellow) and circulatory (white) cells within the Caudal Hematopoietic Tissue (CHT: boxed region in schematic of zebrafish larva, above). Scale bar = 500 μm. (C) Quantification of *drl:mCherry* CTCF levels in treated groups and control groups. (*Tg(drl:CFP-NTR;drl:mCherry)* + 1% DMSO (purple), *Tg(drl:mCherry)* + 10 mM MTZ (light blue); and *Tg(drl:CFP-NTR;drl:mCherry)* + 10 mM MTZ (red) (N = 17-54 larvae). (D) Confocal fluorescent images showing cytoplasmic *drl:CFP-NTR* expression (D’), nuclear *runx1:mCherry* expression (marking HSPCs)(D’’), and merged (D’’’) within the CHT of a 6 dpf zebrafish, with white arrowheads indicating double positive cells. Scale bar = 500 μm. (E) Flow cytometry plots of *runx1:mCherry* and FITC (autofluorescence control) in untreated negative controls (black), *Tg(drl:CFP-NTR;runx1:mCherry)* + 1% DMSO, *Tg(runx1:mCherry)* + 10 mM MTZ; and *Tg(drl:CFP-NTR;runx1:mCherry)* + 10 mM MTZ. (F) Quantification of *runx1:mCherry^+^ %* from (E) flow cytometry experiments in treated and control groups. N = 5-19, 7-10 pooled larvae per sample. Two-way ANOVA with Tukey’s multiple comparisons test was used for all statistical analyses. Plots are individual points for each biological replicate with mean ± standard deviation. ****p-value *≤* 0.0001. See also Figure S5.

The caudal hematopoietic tissue (CHT) is a major site of larval hematopoiesis (equivalent to the mammalian fetal liver) (Murayama, et al., 2006). We detected strong CFP-NTR fluorescent signal in the CHT region in both circulating and stationary cells from 1-8 dpf (Figure S5B). Exposure of *drl:CFP-NTR* embryos to MTZ at 1 dpf to target Prog+ cells resulted in significant mortality and severe edema in larvae by 5 dpf (Figures S5C-D), possibly due to cell death in other *drl*-expressing lineages outside of blood (Mosimann, et al., 2015). In contrast, treatment at 2 dpf, which would target HSCs, showed 100% larval survival and no edema. Control *drl:mCherry* embryos treated with MTZ showed similar mCherry^+^ fluorescence levels and survival as DMSO-treated controls, indicating that MTZ alone does not contribute to significant hematopoietic injury in our system. MTZ treatment of *drl:CFP-NTR* embryos at 2 dpf successfully depleted *drl:mCherry*^+^ cells (Figures 5B-C). We observed significantly fewer circulating and stationary *drl:mCherry^+^* cells in the MTZ-treated experimental group compared to the DMSO- and MTZ-treated controls by 3-4 dpf (or 1-2 days post MTZ treatment [dpM]).

Our *drl*-based NTR/MTZ system enables the study of HSC regeneration within the endogenous developmental environment. We confirmed that HSCs became depleted in the NTR/MTZ system at 2 dpf by using 1) *runx1:mCherry* zebrafish that express mCherry in HSCs (Tamplin, et al., 2015), and 2) *cd41:eGFP* transgenics that express eGFP in HSCs as well as prothrombocytes and differentiated thrombocytes (Lin, et al., 2005). Larval *runx1:mCherry^+^* HSCs express *drl:CFP-NTR* confirmed by confocal microscopy (Figure 5D). Using flow cytometry, we demonstrated a significant depletion of the *runx1:mCherry^+^* HSCs at 3 dpf (1 dpM) and a recovery of this population by 6 dpf (4 dpM) as compared to our control groups (Figures 5E-F). We confirmed the same ablation and regeneration dynamics in *drl:CFP-NTR;cd41:*eGFP double-transgenic embryos (Figures S5E-F). We observed that the nadir of depletion of *cd41:eGFP^+^* cells occurs at 3-4 dpf (1-2 dpM), with some recovery beginning at 5-6 dpf (3-4 dpM). Altogether, these data indicate that embryonic HSCs can regenerate in response to hematopoietic injury.

### HSC depletion minimally impacts myeloid and lymphoid maintenance after hematopoietic injury in early developmental stages

Based on our lineage labeling results, we found that HSCs are not actively contributing to early myeloid and lymphoid larval hematopoiesis (Figures 3-4). We therefore hypothesized that HSC depletion would not affect the maintenance of these lineages during early development as Prog+ predominantly or even exclusively form these lineages. To address this, we examined the levels of *rag2:mCherry^+^* lymphoid cells (Harrold, et al., 2016) and *lysozyme:dsRed^+^* (*lyz*) myeloid cells (Hall, et al., 2007) following HSC depletion. We predicted that if HSCs do play an essential role in early blood production, then their depletion following MTZ treatment would lead to fewer HSC-derived progenitors and ultimately fewer *rag2:mCherry^+^ and lyz:dsRed^+^* mature cells in larvae.

In zebrafish, T lymphocyte progenitors colonize the embryonic thymus by 68 hpf and begin to express *rag2* at 72 hpf (Langenau, et al., 2004; Trede and Zon, 1998). We found no overlap between the lymphocyte marker *rag2:mCherry* and *drl:CFP-NTR* expression in the thymus of 5 dpf zebrafish, confirming T-cells would not be directly affected by MTZ treatment (Figure 6A). After HSC depletion with MTZ at 2 dpf, we monitored *rag2:mCherry* levels in double-transgenic *drl:CFP-NTR;rag2:mCherry* embryos from 3-6 dpf (1-4 dpM) using fluorescence imaging of the thymus and flow cytometry analysis (Figure 6B). By imaging-based analysis, we found MTZ-treated *drl:CFP-NTR^+^;rag2:mCherry^+^* embryos had slightly diminished seeding of the thymus at 3-4 dpf (1-2 dpM) with recovery by 5-6 dpf (3-4 dpM) (Figures 6C-D). However, quantification of T cell frequency by flow cytometry revealed minimal to no decrease of the *rag2:mCherry^+^* cells after HSC depletion (Figures 6E-F). Consistent with our lineage-tracing data, this comparably subtle effect suggests that HSCs have little contribution to larval T lymphocyte production by 6 dpf.

**Figure 6.**
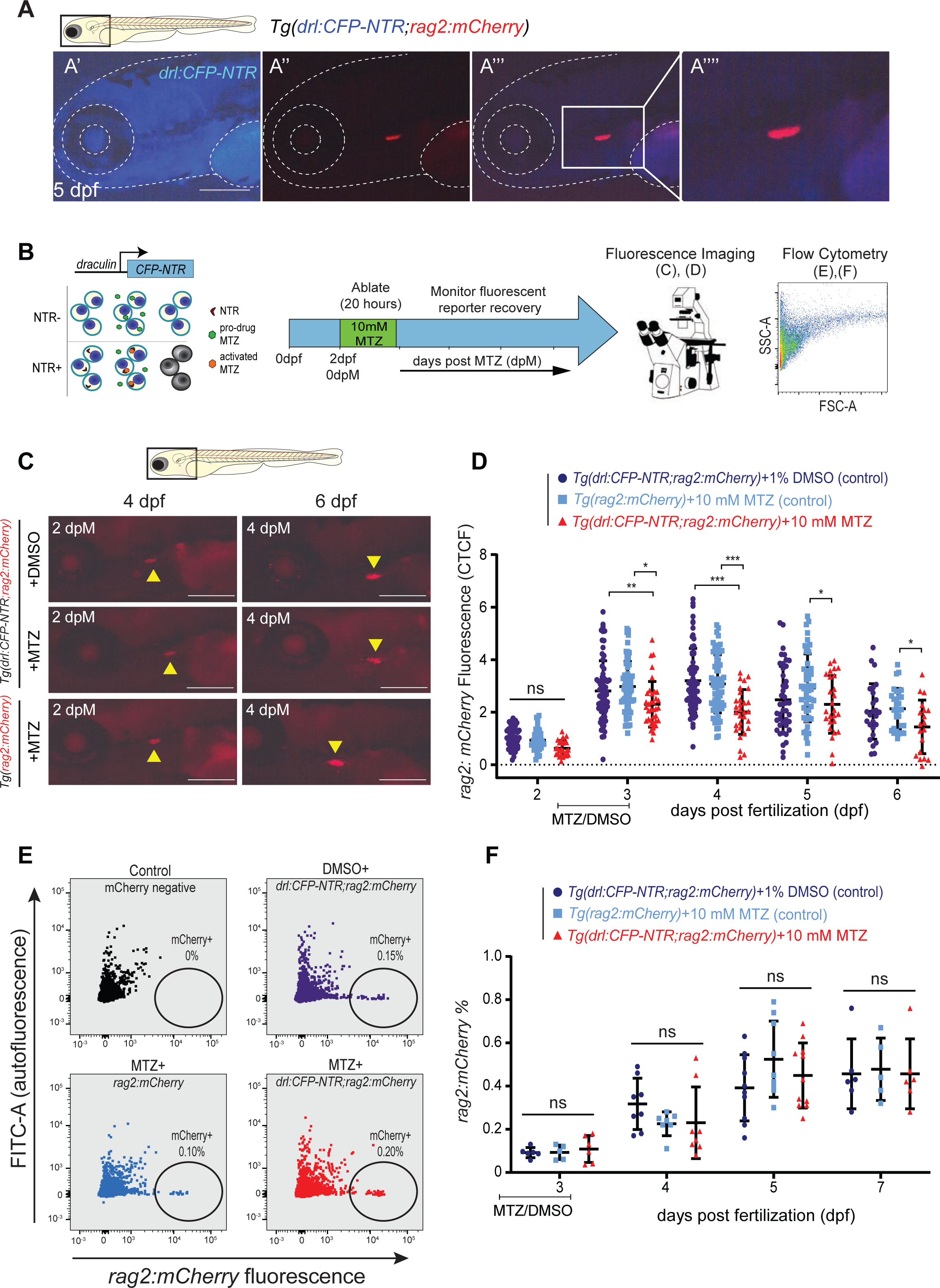
Depletion of HSCs has a negligible impact on T-cell seeding of larval thymus. (A) Fluorescent images of *drl:CFP-NTR* (A’), *rag2:mCherry* (marking T-lymphocytes) (A’’) and merged (A’’’) at 5 dpf in the thymic lymphoid tissue. Scale bar = 500 μm. Merged fluorescent image (A’’’’) shows higher magnification image of (A’’’). (B) Experimental schema of the NTR/MTZ system. (C) Fluorescent images of *Tg(drl:CFP-NTR^+^; rag2:mCherry^+^)* and *Tg(rag2:mCherry^+^)* larvae shown at 4 and 6 dpf (2 and 4 dpM, respectively) after depletion of HSCs using the NTR/MTZ system. Yellow arrowheads indicate *rag2:mCherry*^+^ fluorescent lymphocytes within the thymus. Scale bar = 500 μm. (D) Quantification of *rag2:mCherry* fluorescence CTCF levels in *Tg(drl:CFP-NTR^+^;rag2:mCherry^+^)* and *Tg(rag2:mCherry^+^)* zebrafish treated with 10 mM MTZ or 1% DMSO (N = 19-72 larvae). (E) Flow cytometry plots of *rag2:mCherry* and FITC (autofluorescence control) in untreated negative controls (black); *Tg(drl:CFP-NTR;rag2:mCherry)* + 1% DMSO (purple); *Tg(rag2:mCherry) +* 10 mM MTZ (light blue); and *Tg(drl:CFP-NTR;rag2:mCherry)* + 10 mM MTZ (red). (F) Quantification of *rag2:mCherry%* from (E) flow cytometry experiments in treated and control groups. N = 4-11, 7-10 pooled larvae per sample. Two-way ANOVA with Tukey’s multiple comparisons test was used for all statistical analyses. Plots are individual data points for each biological replicate with mean ± standard deviation. ns not significant, *p-value < 0.05, **p-value *≤* 0.01, ***p-value *≤* 0.001.

The *Tg(lyz:dsRed)* reporter labels lysozyme C-producing cells, mainly granulocytes and some macrophages (Hall, et al., 2007). Similar to lymphoid cells, we found little to no apparent overlap in fluorescence between *drl:CFP-NTR^+^* cells and *lyz:dsRed*^+^ granulocytes (Figure 7A). To measure the effect of HSC depletion on granulocyte levels, we treated double-positive *drl:CFP-NTR;lyz:dsRed* embryos with MTZ at 2 dpf and monitored the *lyz-*expressing cells using fluorescence imaging and flow cytometry (Figure 7B). When imaging the CHT region, we observed a minor decrease of granulocytes with a nadir at 4 dpf (2 dpM) and a full recovery by 6 dpf (4 dpM) as compared to controls (Figures 7C- D). When monitoring the total *lyz:dsRed^+^* cell population using flow cytometry, we found no significant differences between controls and the experimental group (Figures 7E-F). Again, this comparably mild effect of *drl:CFP-NTR^+^* cell depletion on *lyz:dsRed^+^* cells suggests that HSCs do not significantly contribute to larval myelopoiesis up to 7 dpf, consistent with our lineage labeling data.

**Figure 7.**
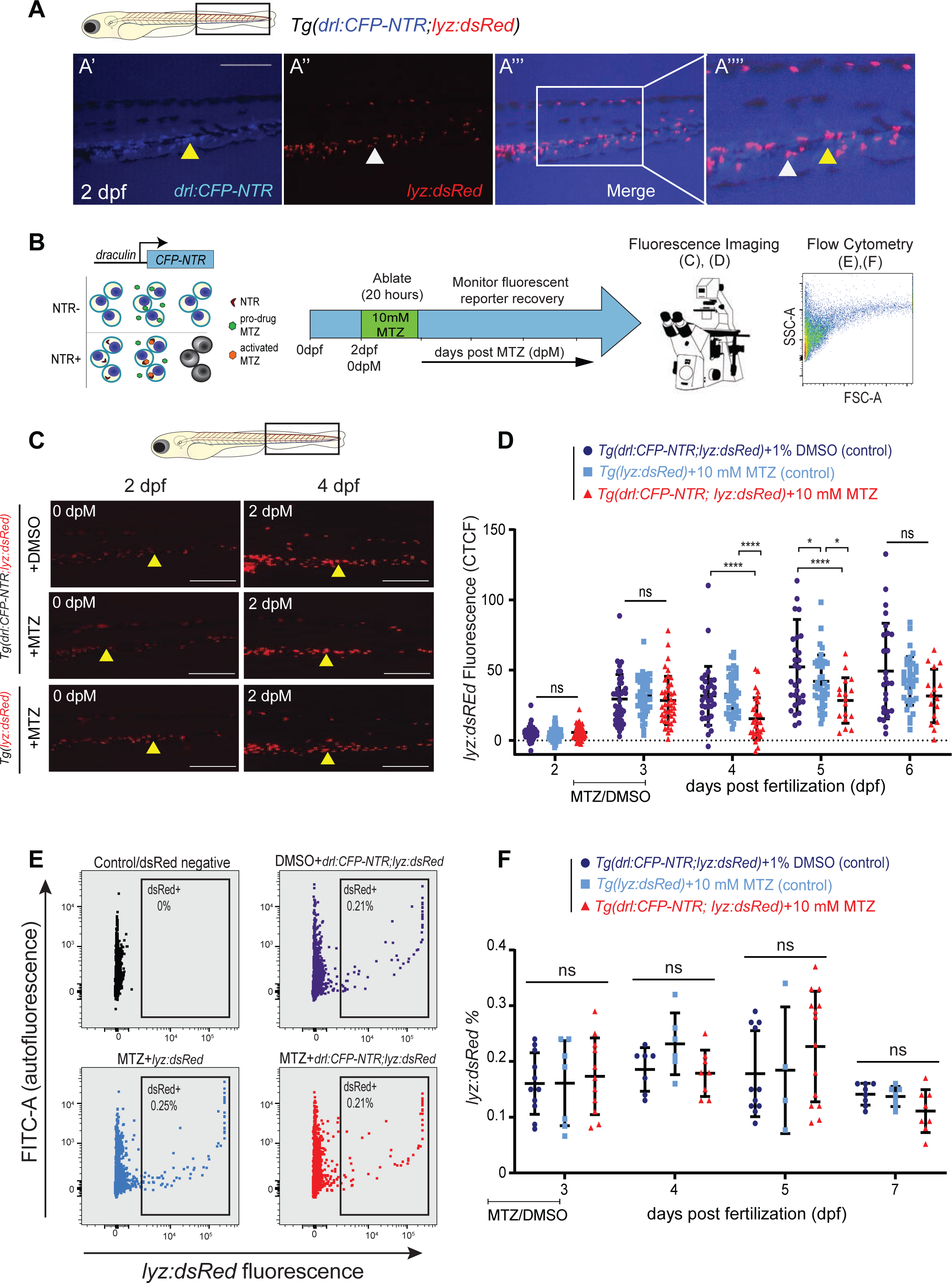
Depletion of HSCs has a negligible impact on granulocyte frequency. (A) Fluorescent images of *drl:CFP-NTR* (A’), *lyz:dsRed* (marking myeloid cells) (A’’) and merged (A’’’) at 2 dpf showing minimal co-expression. Scale bar = 500 μm. White arrow marks *lyz:dsRed* single-positive granulocyte and yellow arrowhead marks *drl:CFP-NTR* single positive cell, 1.75X inset in (A’’’’). (B) Experimental schema of the NTR/MTZ system. (C) Fluorescent images of *Tg(drl:CFP-NTR^+^;lyz:dsRed^+^)* and *Tg(lyz:dsRed^+^)* embryos treated with either 1% DMSO or 10mM MTZ, shown at 2 and 4 dpf (0 and 2 dpM, respectively). Yellow arrowhead showing stationary cells in the CHT region. Scale bar = 500 μm. (D) Quantification of *lyz:dsRed* fluorescence CTCF levels in *Tg(drl:CFP-NTR^+^;lyz:dsRed^+^)* and *Tg(lyz:dsRed+)* embryos treated with 10 mM MTZ or 1% DMSO (n = 16-45). (E) Flow cytometry plots of *lyz:dsRed* and FITC (autofluorescence control) in untreated negative controls (black); *Tg(drl:CFP-NTR;lyz:dsRed)* + 1% DMSO (purple); *Tg(lyz:dsRed)* + 10 mM MTZ (light blue); and *Tg(drl:CFP- NTR;lyz:dsRed)* + 10 mM MTZ (red). (F) Quantification of *lyz:dsRed%* from (E) flow cytometry experiments in treated and control groups (N = 4-14 samples, 7-10 pooled larvae per sample). Two- way ANOVA with Tukey’s multiple comparisons test was used for all statistical analyses. Plots are individual data points for each biological replicate with mean ± standard deviation. Ns not significant, *p-value < 0.05, ****p-value *≤* 0.0001.

These data further confirm that HSCs can regenerate after depletion during embryonic stages and that their depletion has a minor to no impact on larval lymphomyelopoiesis that is driven by Prog+ cells. Combined, our findings document that HSC-independent progenitors, and not HSCs, sustain embryonic hematopoiesis.

## Discussion

A main challenge of studying developmental HSC contribution originates from the difficulty of specifically distinguishing HSCs from embryonic hematopoietic progenitors, as their emergence is spatiotemporally overlapping and commonly used markers are expressed in both developing progenitor and HSC populations (Kasper, et al., 2020; Hadland and Yoshimoto, 2018; McGrath, et al., 2015; Chen, et al., 2011; Godin and Cumano, 2002). Through our scRNA-seq exploration of newly formed HSPCs, we document transcriptional heterogeneity that defines seven distinct HSPC populations including well-documented subsets, such as Pre-HE, HE/HSC, and EryP, and new populations, such as LMP (He, et al., 2020a) and LEP (Kasper, et al., 2020) (Figure 1). This high-resolution transcriptional landscape of developmental HSPCs allowed us to infer differentiation trajectories and cell-type-specific gene enrichment networks relevant to early hematopoiesis. Akin to recent murine scRNA-seq analyses, we have captured pre-HE and HE cells (Zhu, et al., 2020; Zhou, et al., 2016). This pre-HE transition state shows upregulation of vasculature gene expression, while also showing stronger enrichment for hematopoietic development markers. Moreover, we have shown that pre-HE can give rise to cells with distinct development trajectories. However, unlike *Zhu et al*. (2020) who found two different murine hematopoietic waves – an initial lymphomyeloid-biased progenitor followed by pre-HSC precursors (Zhu, et al., 2020) – we discovered at least three distinct trajectories arising from pre-HE cells in zebrafish: HSCs, LyPs, and LEPs (Figure 1D). The granularity with which HSCs and embryonic progenitors are distinguished from each other in our scRNA-seq analysis offers the first comprehensive transcriptomic look into what drives their separation in zebrafish.

That HSCs have a delayed hematopoietic contribution during development indicates a disconnect between the timing of HSC emergence and their demonstrable long-term function. Additionally, our data contrasts the classically held interpretation that HSCs reside at the top of the hematopoietic hierarchy, maintaining lifelong erythroid, myeloid, and lymphoid hematopoiesis (Kobayashi, et al., 2019). Since nascent HSCs can engraft and reconstitute the blood system of a transplanted host, it was posited that they were the source of all blood cells from shortly after their emergence onward (Medvinsky and Dzierzak, 1996). Nonetheless, seminal work in mice has indicated that embryonic hematopoietic progenitors were necessary and sufficient for sustaining the embryo (Chen, et al., 2011). These data suggested that HSCs did not significantly contribute to developmental hematopoiesis, but HSC contribution to differentiated blood lineages was not directly assessed. Although not fully explored in mammals, clonal tracing performed during fetal stages in the mouse has suggested that HSC clones do not robustly contribute to mature lineages until postnatal stages (Busch, et al., 2015). A recent study illustrated that yolk sac-derived progenitors, and not HSCs, sustain erythropoiesis throughout murine embryonic life (Soares-da-Silva, et al., 2021). In developing zebrafish, a T lymphoid progenitor (Tian, et al., 2017) and an LMP population (He, et al., 2020a) were also recently identified to arise in development and represent distinct HSC-independent progenitors. Consistent with these studies, we demonstrate that embryonic hematopoietic progenitors in zebrafish sustain early developmental hematopoiesis while HSCs do not detectably contribute to larval lymphomyelopoiesis (Figures 3-4). Combined, these findings in zebrafish and murine models indicate a strong conserved differentiation latency of nascent HSCs spanning all hematopoietic lineages.

The observed differentiation latency itself could also denote the nascency or immaturity of HSCs. In adult organisms, injury-induced regeneration assays are often used to challenge HSCs to illuminate function. These approaches are mainly based on myeloablative injuries, such as chemotherapy or irradiation that target proliferating cells. Developing embryos are highly proliferative, thus use of such treatments impair overall development and organ formation. To investigate the potential for HSC regeneration and their temporal lineage contribution during development *in situ*, we here established an HSC-selective injury model based on transgenic *NTR* driven by the *drl* regulatory elements that permitted an assessment of HSC stress response during zebrafish ontogeny without transplantation. We delineated that HSCs regenerate following their depletion, but that this hematopoietic injury had a negligible impact on developmental lymphomyelopoiesis which is instead dependent on earlier hematopoietic progenitor cells (Figures 5-7). The demonstration of HSC regeneration during development is consistent with studies in zebrafish and mice that show self-renewal capacity of embryonic HSCs upon transplantation (Tamplin, et al., 2015; Ema and Nakauchi, 2000; Müller, et al., 1994). Future studies using our developmental regeneration assay have the potential to provide novel insights into the spatio-temporal dynamics of HSC self-renewal and differentiation properties.

Together, our findings demonstrate that after their emergence, HSCs display differentiation latency but active self-renewal. Our data illustrate that the embryonic progenitors, not HSCs, sustain early developmental hematopoiesis. Understanding how and where HSCs might acquire the ability to sense and regenerate a challenged or damaged blood system during development will help guide improvements to generate functional HSCs from renewable pluripotent stem cells. Additionally, studying HSC self-renewal and differentiation during ontogeny may help us delineate the factors that promote HSC expansion without loss of other functions.

## Acknowledgements

This work was funded by American Cancer Society RSG-129527-DDC, DOD BM180109, NIH 1R01DK121738-01A1 and the Edward P. Evans Foundation (to TVB), NIH T32GM007288 and F31HL152562 (to BAU), Einstein SOARS program (to SSH), the American Australian Association Sir Rupert Murdoch Postdoctoral Fellowship (to KSP), The Einstein Training Program in Stem Cell Research funded by the Empire State Stem Cell Fund through New York State Department of Health Contract C30292GG (to KSP and AN), and funds from The University of Colorado School of Medicine and the Children’s Hospital Colorado Foundation (to CM). Instrumental feedback to the growth of this project was provided by Britta Will, Kira Gritsman, Ulrich Steidl, and Sofia De Oliveira. We would like to thank several Einstein Core Facilities supported by NIH P30CA013330, including Flow Cytometry, Genomics, and Computational Genomics Facilities. For their assistance with cell sorting and flow cytometry analysis, we thank the Flow Cytometry core members Jinghang Zhang, Yu (Joey) Zhang, Fnu (Aodeng) Aodengtuya, and Ming Liu. Additionally, we would like to thank the Einstein Genomics Core and David Reynolds for their assistance with 10X Genomics scRNA pre-processing and library preparation. For their computational support, thank you to Tobias Schraink and the Einstein Computational Genomics Core, Dr. Robert Dubin. For their animal support, we thank the Einstein Zebrafish core members Clinton de Paolo and Spartak Kalinin. Lastly, we greatly thank the Hadland lab and its members, especially Dr. Brandon Hadland and Tessa Dignum, for their guidance with scRNAseq analysis and providing code.

## Author Contributions

Conceptualization, TVB, BAU, SSH; Methodology, TVB, BAU, SSH, KSP, AN, AL, MM, SP, JF, MFN, CM; Investigation, TVB, BAU, SSH, AN, JF; Writing—Original Draft, TVB, BAU, SSH; Writing—Review & Editing, TVB, BAU, KSP, CM; Funding Acquisition, TVB, BAU; Supervision, TVB.

## Declaration of Interests

The authors declare no competing interests.

## STAR METHODS

### KEY RESOURCES TABLE

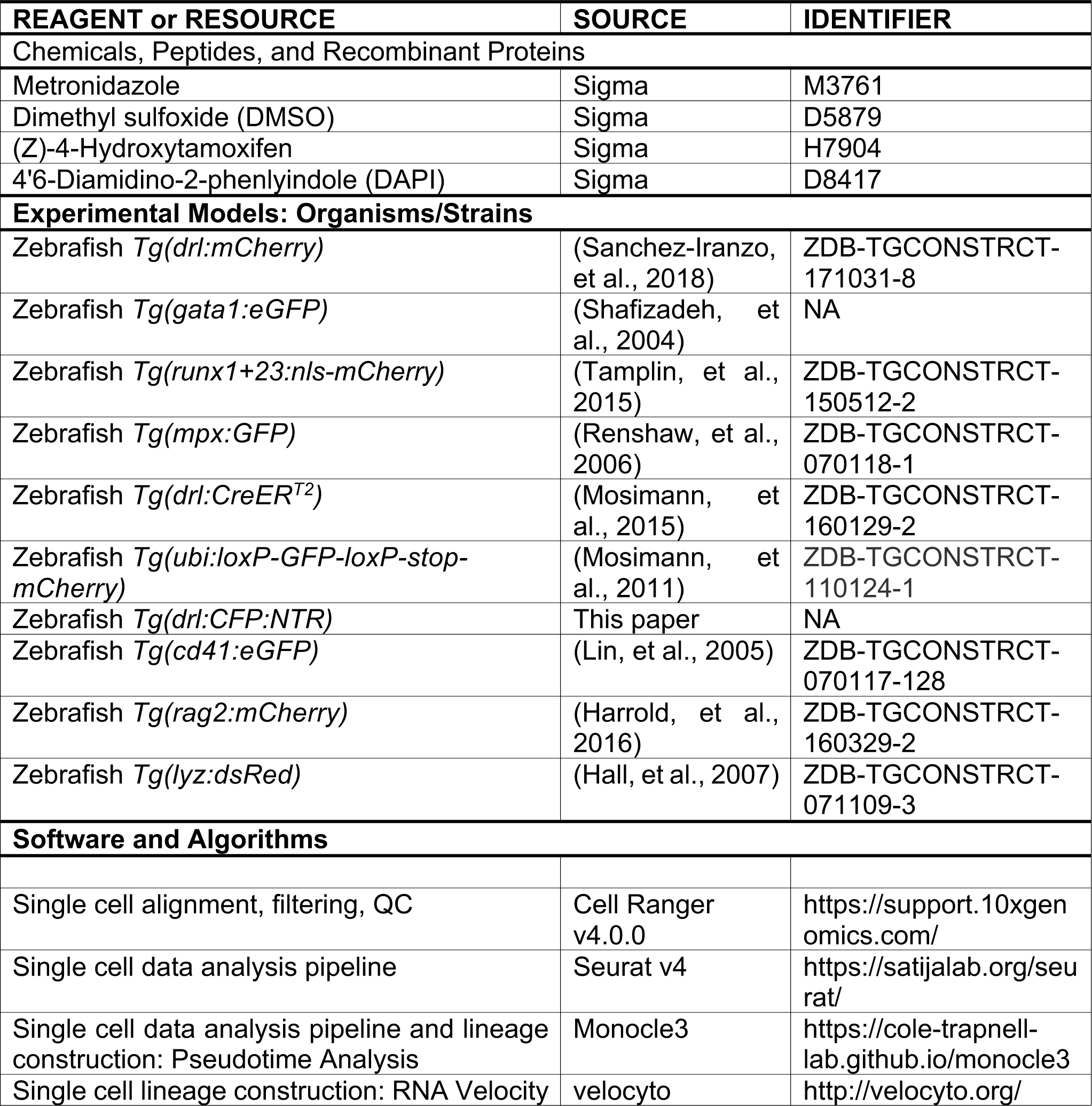
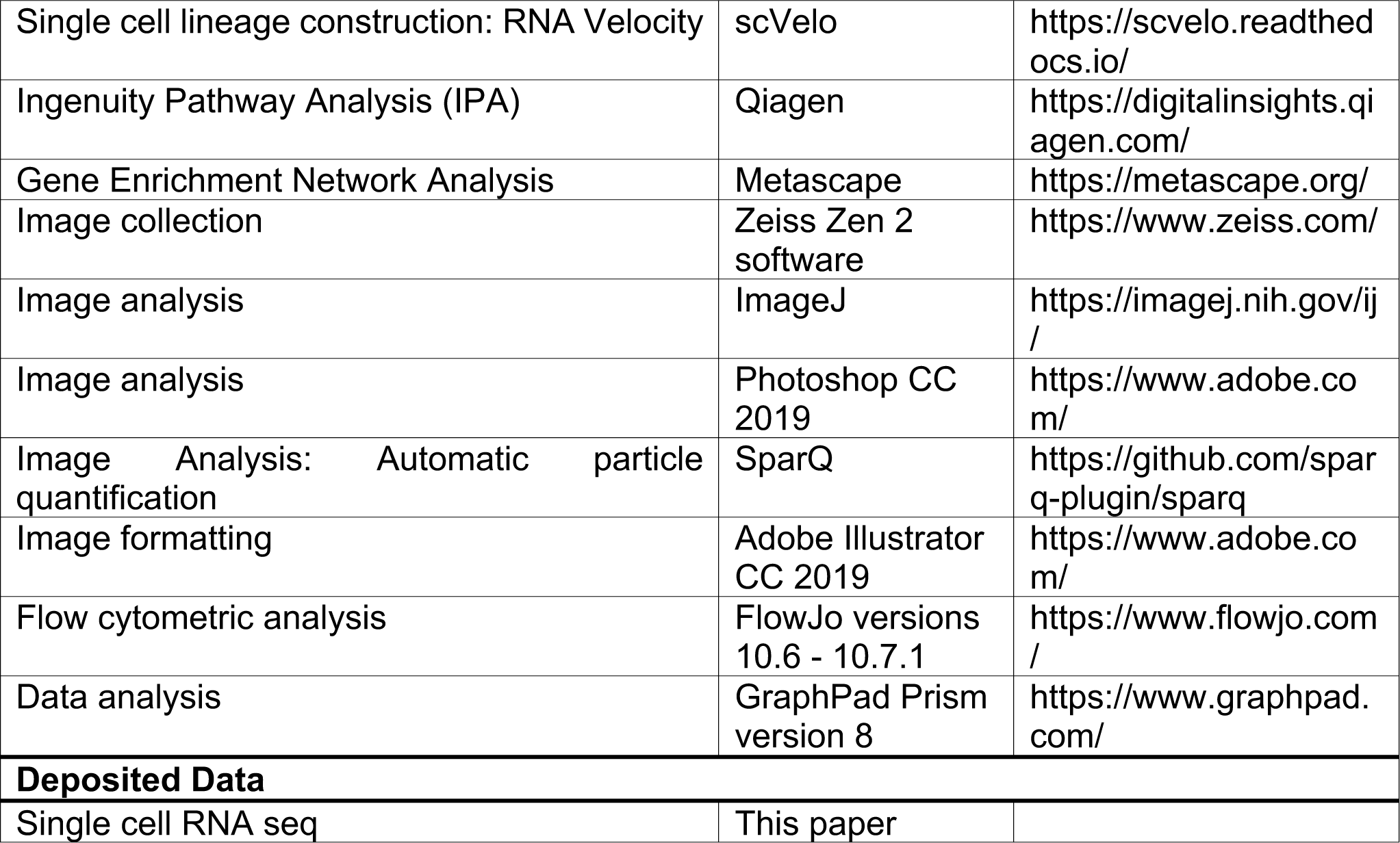

### LEAD CONTACT AND MATERIAL AVAILABILITY

Further information and requests for scripts, resources, and reagents should be directed to and will be fulfilled by Lead Contact, Teresa V. Bowman.

### EXPERIMENTAL MODEL AND SUBJECT DETAILS

#### Zebrafish husbandry

Zebrafish were bred and maintained as described previously (Lawrence, 2011). All fish were maintained according to Institutional Animal Care and Use Committee (IACUC)-approved protocols in accordance with the Albert Einstein College of Medicine research guidelines.

#### Transgenic lines

Transgenic lines used in this study included *Tg(drl:mCherry)* (Sanchez-Iranzo, et al., 2018)*, Tg(gata1:eGFP)* (Shafizadeh, et al., 2004)*, Tg(runx1*+*23:nls-mCherry*) [shortened to *Tg(runx1:mCherry)*] (Tamplin, et al., 2015), *Tg(mpx:GFP)* (Renshaw, et al., 2006)*, Tg(cd41:eGFP)* (Lin, et al., 2005), *Tg(rag2:mCherry)* (Harrold, et al., 2016), and *Tg(lyz:dsRed)* (Hall, et al., 2007). For the lineage tracing experiments, we used *Tg(drl:creER^T2^)* (Mosimann, et al., 2015)*, Tg(ubi:loxP-GFP-loxP-stop-mCherry)* [also known as *Tg(ubi:Switch)*] (Mosimann, et al., 2011), and *Tg(mpx:GFP;ubi:Switch),* which are the progeny of *Tg(mpx:GFP)(Renshaw, et al., 2006)* crossed to *Tg(ubi:switch)*.

The *drl:CFP-Nitroreductase (NTR)* construct was generated using multisite gateway cloning. The *drl* regulatory elements in pCM296 as *pENTR5’* vector (Mosimann, et al., 2015), *CFP-NTR* construct (Curado, et al., 2007), *α*-*crystallin:YFP*-carrying *pDestCY* backbone *pCM326* (Mosimann, et al., 2015) have been previously described. The construct and *Tol2* RNA (Kwan, et al., 2007) were injected into one-cell stage embryos, which were then screened for correct CFP expression. Positive embryos were reared to adulthood and screened for positive *F0* founders. Single-insertion transgenic strains with persistent expression of *drl:CFP-NTR* in only blood (and not dominantly expressed in the heart or endothelium) were specifically selected to prevent toxicity from injuring these essential tissues where *drl*-based transgenics can also be actively expressed (Prummel, et al., 2019; Mosimann, et al., 2015). All experiments were confirmed with at least two independent transgenic lines.

#### Embryo/larval fluorescence-activated cell sorting

Flow cytometry protocol was used to prepare *Tg(drl:mCherry;gata1:GFP) (*30 and 52 hpf) and *runx1:mCherry* (52 hpf) in replicate for fluorescent cell sorting prior to 10X Genomics scRNA-seq. For each group, 200-300 embryos were manually dissociated using a sterile razor blade. The material was then resuspended in 1 mL of 1x Dulbeccos-PBS (D-PBS) (Life Technologies) and digested with a 1:65 dilution of 5 mg/mL Liberase (Roche) for 6 minutes in a 37°C water bath. The reaction was stopped with the addition of 5% fetal bovine serum (FBS) (Life Technologies). The triturated suspension was then filtered twice through a 40 μm cell strainer (Falcon), pelleted by centrifugation, and resuspended in 2 mL of FACS buffer (0.9x D-PBS, 5% FBS, 1% Pen/Strep (Life Technologies)). DAPI (4’,6- diamidino-2-phenylindole) was added to a final concentration of 1 μg/mL to exclude dead cells from the analysis. Gates for background signal were drawn using cells from non-fluorescent embryos (Figures S1A-B). For *Tg(drl:mCherry;gata1:GFP)*, we sorted mCherry-positive and GFP-negative cells at 1 and 2 dpf (28-30 and 50-52 hpf, respectively) in replicate to enrich for HSPCs and decrease the erythroid *gata1* signal. For *Tg(runx1:mCherry)*, we sorted for mCherry-positive cells at 2 dpf (50-52 hpf). Sorting was performed on a Becton Dickinson FACSAria which was equipped with a 100-micron nozzle, run at pressure 20 psi with flow rates less than 3,000 events/second. Sorted cells were collected into microcentrifuge tubes containing 500 μL IMDM + 10% FBS at 4°C.

#### scRNA-Seq

Sorted cells were processed for library preparation using the 10X Genomics Chromium Single Cell 3’ Reagent Kit v3.1.0, performed by the Einstein Genetics Genomics Core. Libraries were then sent to Genewiz for sequencing. Initial sample quality control (QC) involved assessing library size on the Agilent TapeStation Analysis Software 3.2 (Agilent Technologies, Inc 2019), concentration measurement on Qubit dsRNA Assay, and final quantitation by qPCR. Indexed libraries were pooled and sequenced on an Illumina HiSeq paired-end 2 x 150 bp read length, single index with 5% PhiX spike-ins.

### scRNA-seq data analysis

#### Preprocessing of scRNA-seq data

Sample alignment, filtering, barcode counting, and unique molecular identifier counting (UMI) were performed with 10x Genomics Cell Ranger 3.1.0 pipeline. Samples were aligned to a custom zebrafish reference genome (*Danio rerio* GRCz11/danRer11) that included *mcherry* and *gfp* sequences, built using the Cell Ranger cellranger *mkref* pipeline. We sequenced 25,574 cells of which 25,248 cells passed quality control using Cell Ranger, including 4,469 cells at 1 dpf (30 hpf) and 20,779 cells at 2 dpf (52 hpf). On average, 1804 and 676 genes were detected per cell at 1 and 2 dpf embryos, respectively (Supplemental Table 1). Cells that passed quality control metrics as defined by the Cell Ranger pipeline were passed into downstream clustering analysis.

#### Dimension reduction, unsupervised clustering, and cell type identification

Seurat (version 4.0)(Hao, et al., 2020) and Monocle3 (Cao, et al., 2019) were used to analyze merged *drl:mCherry*^+^;*gata1:GFP^-^* samples (replicates for 1 dpf and replicates for 2 dpf). For Seurat, we preprocessed (nFeature_RNA 200-4500 and mitochondrial percentage <5%), normalized, and selected 5,000 most variable genes to feed into dimension reduction by principal component analysis PCA. Then, the top 50 principal components were selected for uniform manifold approximation and projection (UMAP) reduction and clustering. The number of clusters is dependent on parameter resolution (0.5), from which we resolved 12 cluster groups. SeuratWrappers were used to convert into a Monocle3 cell dataset object, which we used for trajectory and top marker analysis. For Monocle3, we selected the top 50 dimensions for preprocessing using PCA method, and for dimensional reduction using UMAP with a resolution of 7e-5, by which the number of cluster groups were 10. Batch-correction (Haghverdi, et al., 2018) was conducted prior to UMAP dimensional reduction by matching mutual nearest neighbors. For the overlap of *drl:mCherry*^+^;*gata1:GFP*^-^ and *runx1:mCherry^+^* samples at 2 dpf, the above protocol was also followed.

Cell types were assigned by key marker gene expression, as defined by literature search. Top marker analysis was conducted based on cell type assignments, up to 200 genes were chosen per group (criteria: fraction expressed in > 10% of cells and a test p value < 0.00005). Top markers derived for each cell type using Monocle3, as described above, were then used for pathway enrichment analysis with Metascape (Zhou, et al., 2019) and Ingenuity Pathway Analysis (IPA, QIAGEN).

#### HE/HSC zebrafish and murine comparison

In Seurat v4.0, we conducted differential expression (DE) analysis for upregulated genes (p > 3.89E-6, average log2FC > 0.693) in the HE/HSC cluster (c6) using non-parametric Wilcoxon rank sum test. This HE/HSC list of DE genes was compared to publicly available murine HE (Hou, et al., 2020; Zhu, et al., 2020) and pre-HSC/HSC (Vink, et al., 2020; Zhu, et al., 2020; Zhou, et al., 2016) datasets. We converted the gene lists from these studies into their zebrafish homologue counterparts with online tool Ensembl Biomart. Code to derive gene set scores were provided by the Hadland lab. Single cell gene set scores were calculated as the log-transformed sum of the size factor-normalized expression for each gene in publicly available murine datasets: pre-HE (Zhu, et al., 2020) and HE/HSC (Hou, et al., 2020; Vink, et al., 2020; Zhu, et al., 2020; Zhou, et al., 2016).

#### Differential trajectories

Cell Ranger output files for *drl:mCherry*^+^;*gata1:GFP*^-^ samples (replicates for 1 dpf and replicates for 2 dpf) were read into velocyto (La Manno, et al., 2018) for the creation of .loom formatted files containing spliced, unspliced, and ambiguous counts. These reformatted data were merged and reprocessed in Seurat. Preprocessing, dimensional reduction with UMAP, and cell type identification were conducted as described above for the Seurat pipeline. Using this new cluster projection, we were able to identify the same cell types as before. SeuratWrappers and publicly available tutorials on Github helped to create inferred lineage differentiation trajectories using RNA velocity (scVelo) and pseudotime analysis (Monocle3).

#### Lineage Tracing with *Tg(ubi:Switch)*

For lineage tracing experiments using *Tg(drl:creER^T2^;ubi:Switch)* or *Tg(drl:creER^T2^;mpx:GFP;ubi:Switch)*, embryos at 30 or 54 hpf were transferred to a petri dish (150 mm x 15 mm) at a density of 40-50 embryos per plate. Embryo buffer was replaced with E3 embryo buffer (5 mM NaCl, 0.17 mM KCl, 0.25 mM CaCl2, and 0.15 mM MgSO4) containing 12 µM (Z)-4- Hydroxytamoxifen (4-OHT) (Sigma, H7904) or 0.05% (v/v) ethanol (EtOH) as a vehicle control (Henninger, et al., 2017). Zebrafish were treated with 4-OHT or EtOH for 20 hours (30-50 hpf or 54-74 hpf) in a 28°C incubator. After treatment, embryos were washed three times with fresh embryo buffer and placed back in a 28°C incubator. From 5-16 dpf, larvae were transferred to the fish facility nursery where they were kept in fish water supplemented with methylene blue with E3 embryo medium at room temperature and fed paramecia twice daily.

#### Nitroreductase (NTR)/Metronidazole (MTZ) Assay

For assessing the impact of *drl*-cell depletion on the blood system and to determine co-expression of *drl:CFP-NTR* will specific cell types*, Tg(drl:CFP-NTR)* adult zebrafish were crossed to cell type-specific mCherry fluorescent transgenic lines or AB WT fish. At 24 hpf, the embryos were scored for both CFP and mCherry fluorescence. As the mCherry fluorescence is brighter than CFP, mCherry fluorescence was used to monitor the impact of *drl* cell depletion on each lineage. Cell type specific transgenic lines used were *drl-Tg(drl:mCherry)* (Sanchez-Iranzo, et al., 2018), HSPC/thrombocytes-*Tg(cd41:eGFP)* (Lin, et al., 2005), HSPC-*Tg(runx1:mCherry*) (Tamplin, et al., 2015), granulocytes- *Tg(lyz:dsRed)* (Hall, et al., 2007), and T-cells- *Tg(rag2:mCherry)* (Harrold, et al., 2016).

At 54 hpf (0 dpM), embryos were treated as such: *Tg(drl:CFP-NTR)-*positive embryos were treated with either 1% (v/v) DMSO or 10 mM MTZ (Sigma, M3761), and *Tg(drl:CFP-NTR)^-^* embryos were treated with 10 mM MTZ. Dilutions of DMSO or 1 M MTZ stock were made with E3 embryo medium. Embryos were then placed into 24-well plates (n=15/well) with 2 mL of solution per well or in petri dishes (150 mm x 15 mm, n=40-50) with 25 mL of solution per plate and treated for 20 hours overnight in a 28°C incubator. Light exposure was avoided by wrapping the plates in foil as MTZ is light sensitive. Embryos were then washed twice with E3 water. Zebrafish were analyzed via fluorescence imaging and flow cytometry methods.

### Fluorescent imaging

Live zebrafish embryos (54 hpf – 8 dpf) were anesthetized with 0.01% tricaine (Fisher), then mounted in 4-6% (wt/vol) methylcellulose in 35-mm imaging dishes (MatTek) as described previously (Renaud, et al., 2011). Fluorescent imaging of *Tg(drl:creER^T2^;ubi:Switch), Tg(drl:CFP-NTR;drl:mCherry), Tg(drl:CFP-NTR;cd41:GFP)*, *Tg(drl:CFP-NTR;lyz:dsRed),* and *Tg(drl:CFP-NTR;rag2:mCherry)* were performed with Zeiss Discovery.V8 and Zeiss Axio Observer A1 Inverted microscope with an AxioCam HRc Zeiss camera and Zeiss Zen 2 or 2.5 software. Fluorescence was detected with cyan fluorescent protein (CFP), mCherry, Texas Red (for dsRed lines), and green fluorescent protein (GFP) filters.

A Zeiss Live DuoScan confocal microscope with AIM 4.2 Software was used to visualize co-expression in *Tg(drl:CFP-NTR^+^;runx1:mCherry^+^)* embryos using 405nm and 561nm excitation wavelength. Embryos (6 dpf) were anesthetized with 0.01% tricaine (Fisher), oriented in a drop of 3% wt/vol methylcellulose, then mounted in 1% agarose in 35-mm imaging dishes (MatTek) as described previously (Renaud, et al., 2011).

### Fluorescence Microscopy Image Analysis

Fluorescence microscopy images of the raw data were analyzed using FIJI (Schindelin, et al., 2012). On each image, a region of interest (the fluorescent tissue within the CHT or thymus) was selected. Corrected Total Cell Fluorescence (CTCF) was calculated as Integrated density of region of interest - (Area of region of interest X Mean fluorescence of 3 to 6 different background regions). CTCF values of experimental embryos were normalized to that of the controls by dividing the CTCF value of an individual embryo’s image by the average CTCF value of all the images from the control group. All data were then statistically analyzed (see below). Edematous embryos were not included in the analysis.

To determine the threshold for mCherry+ thymi in the lineage tracing analysis, 7 scientists blindly scored for mCherry+ thymi in 34 images of 5 and 10 dpf *Tg(drl:creER^T2^;ubi:Switch)* zebrafish previously treated with 4-OHT from 30-50 hpf or 54-74 hpf. The frequency at which each image was scored positive and negative among the scientists was calculated. Images that were scored positive more than 50% of the time were deemed to have a detectable mCherry^+^ thymus. We calculated the sensitivity and specificity of using CTCF cut-offs by this visual detection standard. A true positive (TP) was an image positively scored and having a CTCF above the cut-off; a false negative (FN) was an image positively scored but having a CTCF below the cut-off; a true negative (TN) was an image negatively scored and having a CTCF below the cut-off, and a false positive (FP) is an image negatively scored and having a CTCF above the cut-off. By plotting a receiving operative characteristics (ROC) curve, we determined a CTCF threshold (7.5) that would give us both an optimal sensitivity (TP/[TP+FN]) and specificity (TN/[TN+FP]) (Figure S6B).

The SparQ (Streamlined Particle Quantification)(Mesquita, et al., 2020) software was used to automate the quantification of the fluorescence particles of *Tg(drl:CFP-NTR; cd41:GFP)* zebrafish.

### Flow cytometry protocol and analysis

Flow cytometry analysis was conducted for *Tg(drl:creER^T2^;mpx:GFP;ubi:Switch)* as part of the lineage tracing assay from 2-16 dpf and for *runx1:mCherry, lyz:dsRed,* and *rag2:mCherry* as part of the NTR/MTZ regeneration assay from 2-9 dpf. For each group, 7-10 embryos were anesthetized with 0.01% tricaine (Fisher) and then dissociated using a sterile razor blade. The material was then resuspended in 600 μL of 1x Dulbeccos-PBS (D-PBS) (Life Technologies) and digested with a 1:65 dilution of 5 mg/mL Liberase (Roche) for 10-15 minutes in 37°C water bath. The reaction was stopped with the addition of 5% fetal bovine serum (FBS) (Life Technologies). The triturated suspension was then filtered through a 40 μm cell strainer (Falcon), pelleted by centrifugation, and resuspended in 300 μL of FACs buffer (0.9x D-PBS, 5% FBS, 1% Penn/Strep (Life Technologies)). DAPI was added to a final concentration of 1 μg/mL to exclude dead cells from the analysis. Gates for background signal were drawn using cells from non-fluorescent embryos.

For adult kidney marrow analysis, kidneys from 3-4 months post fertilization zebrafish were dissected, resuspended in 500 μL of FACS buffer supplemented with 1 μg/mL DAPI, and filtered through a 40-μm cell strainer (Falcon). Forward and side scatter parameters were used to resolve the major blood cell lineages of erythroid, myeloid, lymphoid, and precursor, as previously described (Traver, et al., 2003).

All samples were analyzed at the Flow Cytometry Core Facility at the Albert Einstein College of Medicine using an LSR II flow cytometer (BD Biosciences) and data was processed using FlowJo Software (versions 10.6 - 10.7.1).

### Statistical Analysis

Kaplan-Meier curve analysis and two-way ANOVA with Sidak’s and Tukey’s multiple comparisons tests were performed using GraphPad Prism (version 8). Experiments were performed with a minimum of three independent replicates. Error bars indicate standard deviation from mean, or as specified. ns, not significant; * p<0.05; ** p<0.01; *** p<0.001; **** p<0.0001.

## Supplementary Figures and Tables

Figure S1, related to Figure 1. Supplemental scRNA-seq analysis.

Figure S2, related to Figure 2. Supplemental *Switch* mCherry imaging analysis for Prog+ and HSC populations.

Figure S3, related to Figure 3. Supplemental lymphoid lineage tracing quantification for Prog+ and HSC populations.

Figure S4, related to Figure 4. Supplemental myeloid lineage tracing gating strategy for Prog+ and HSC populations.

Figure S5, related to Figure 5. Supplemental information for the establishment of *in situ* HSC regeneration assay using NTR/MTZ system in zebrafish.

Supplemental Table 1. Summary Statistics from 10X Genomics Cell Ranger from *drl:mCherry+gata1:GFP* at 1 and 2 dpf and runx1:mCherry at 2 dpf

Supplemental Table 2. Top markers (Monocle3) and gene ontology analysis (Metascape) for the different HSPCs in developmental hematopoiesis

Supplemental Table 3. Zebrafish HE/HSC cluster comparison with murine HE and HSC signatures

Supplemental Table 4. Quantification for the number of cells found within each cell type category (Pre-HE, HE/HSC, LyP, LMP-M, LMP-G, LEP, and EryP)

## Notes

### Competing Interest Statement

The authors have declared no competing interest.

