## Supplemental Figures for "Definitive Hematopoietic Stem Cells Minimally Contribute to Embryonic Hematopoiesis"

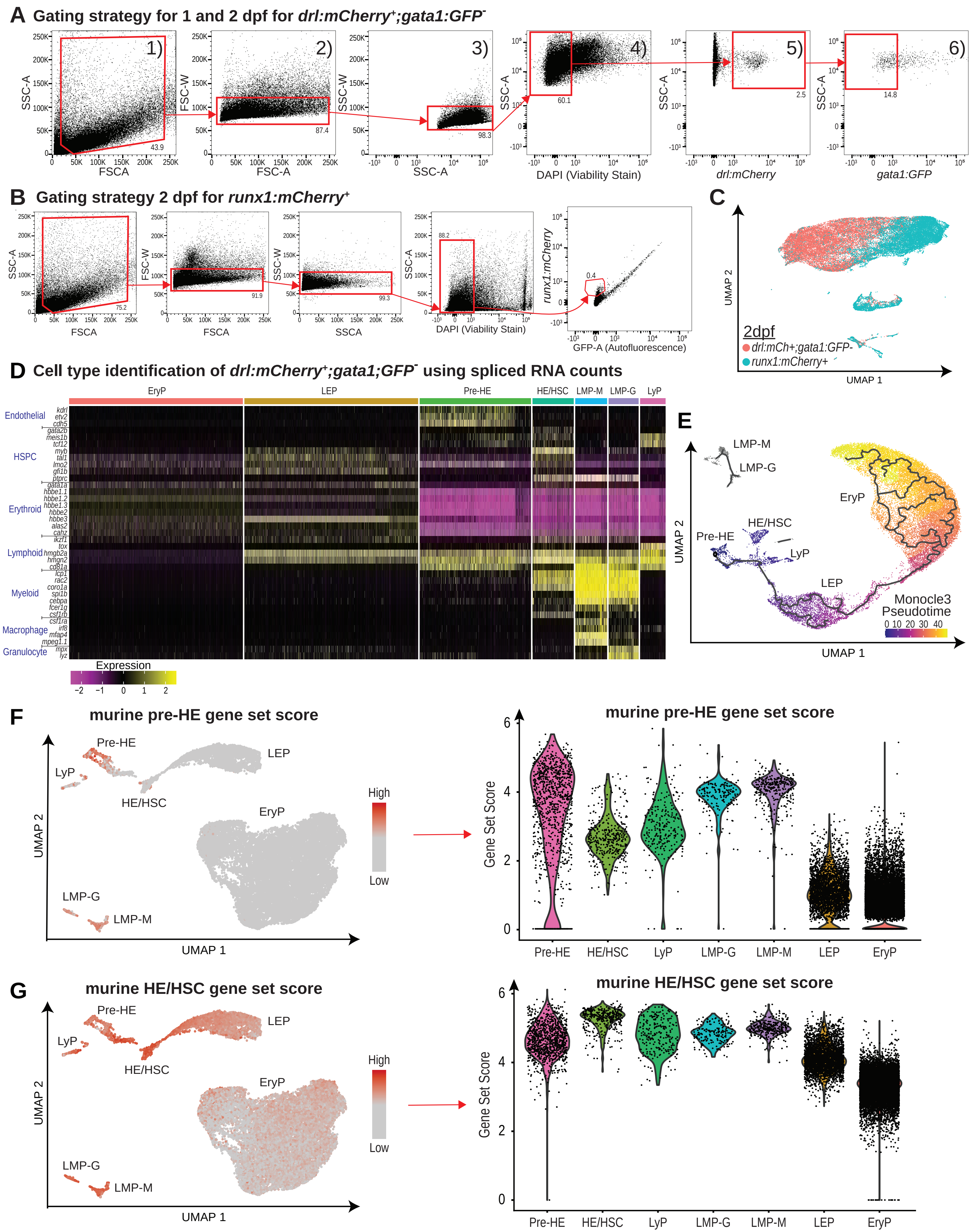


**Supplemental Figure 1.**

**Figure S1, related to Figure 1**. Supplemental scRNA-seq analysis. (A) Flow cytometry gating strategy for *drl:mCherry*$^{+}$ *gata1:GFP*$^{-}$ cells to enrich for HSPCs at 1 and 2 dpf: 1) Forward scatter (FSC)-A vs side scatter (SSC)-A to exclude debris; 2) FSC-A vs FSC-W to select for single cells; 3) SSC-A vs SSC-W to select for single cells; 4) DAPI vs SSC-A to select for DAPI-negative viable cells; 5) mCherry-A vs SSC-A to select for *drl:mCherry*$^{+}$ cells; 6) GFP-A vs SSC-A to select for *gata1:GFP*-negative cells. (B) Flow cytometry gating strategy for *runx1:mCherry*-positive cells to enrich for HSPCs at 2 dpf using similar gating steps 1-4 as in (A). To select for *runx1:mCherry*$^{+}$ cells, GFP-A (autofluorescence) vs mCherry-A to select for *runx1:mCherry*$^{+}$ cells. Non-fluorescent wild-type controls were used to set gates for fluorescent protein expression (not shown). (C) UMAP dimensional reduction of *drl:mCherry*$^{+}$*;gata1:GFP*$^{-}$ and *runx1:mCherry*$^{+}$ cells at 2 dpf. (D) Cell type re-identification for UMAP clustering in Figure 1D, S1E. Expression bars for heatmaps: scaling is performed per gene where mean is close to 0 and standard deviation of 1. (E) Monocle3 pseudotime trajectory overlaid over UMAP dimensional reduction of *drl:mCherry*$^{+}$*; gata1:GFP*$^{-}$ cells at 1 and 2 dpf. Pre-HE identified as the start of pseudotime, or least differentiated, with EryP being at the end of the trajectory, or most differentiated cell state. (F-G) Left: Heatmap of gene-set scores for previously reported murine signature gene sets defining (F) pre-HE (Zhu, et al., 2020) and (G) HE//HSC (Hou, et al., 2020; Vink, et al., 2020; Zhu, et al., 2020; Zhou, et al., 2016). Right: Violin plots of gene set scores by cell type.

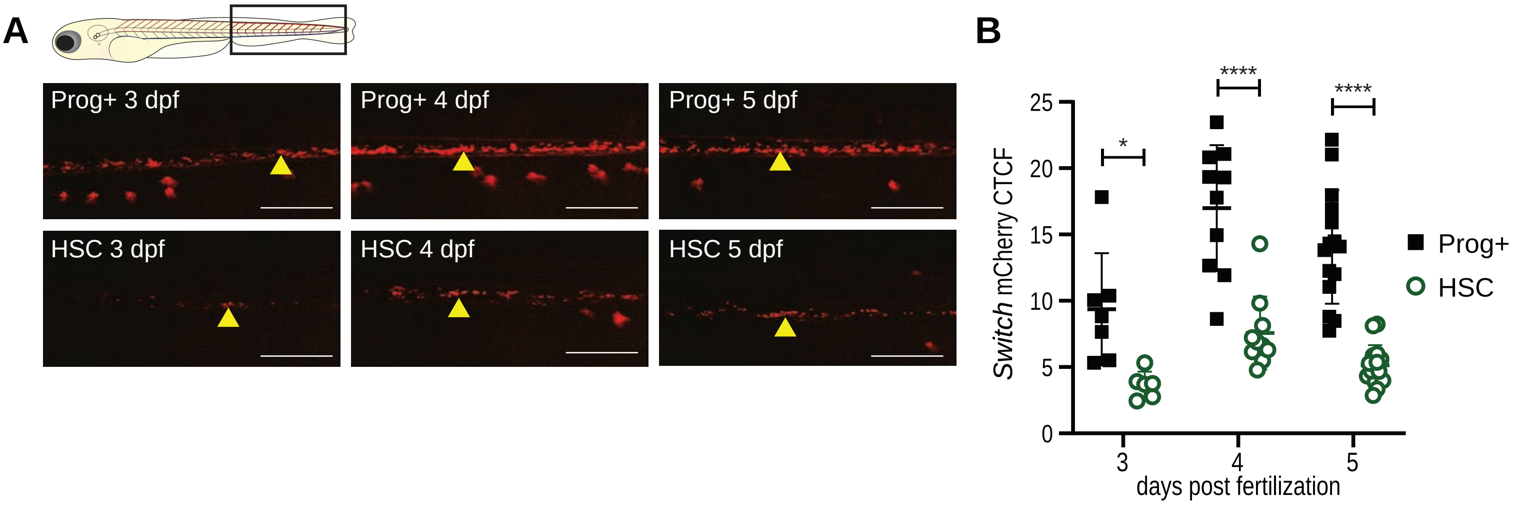


**Supplemental Figure 2.**

**Figure S2, related to Figure 2**. Supplemental *Switch* mCherry imaging analysis for Prog+ and HSC populations. (A) Fluorescent images of *Tg(drl:creER$^{T2}$;ubi:Switch)* embryos for Prog+ (top panel) and HSC (bottom panel) mCherry-labeled populations at 3, 4, and 5 dpf. Yellow arrowhead indicates stationary HSPCs in the caudal hematopoietic tissue (CHT). Scale bar = 500 μm. Representative images shown, quantification in B. (B) Quantification of mCherry$^{+}$ CHT fluorescence from Prog+ and HSC lineage tracing cohorts. Mean ± standard deviation of the *Switch* mCherry$^{+}$ corrected total cell fluorescence (CTCF) at each time point is shown. Two-way ANOVA with Sidak's multiple comparison (N = 7-15 larvae/day). *p-value < 0.05, ****p-value ≤ 0.0001.

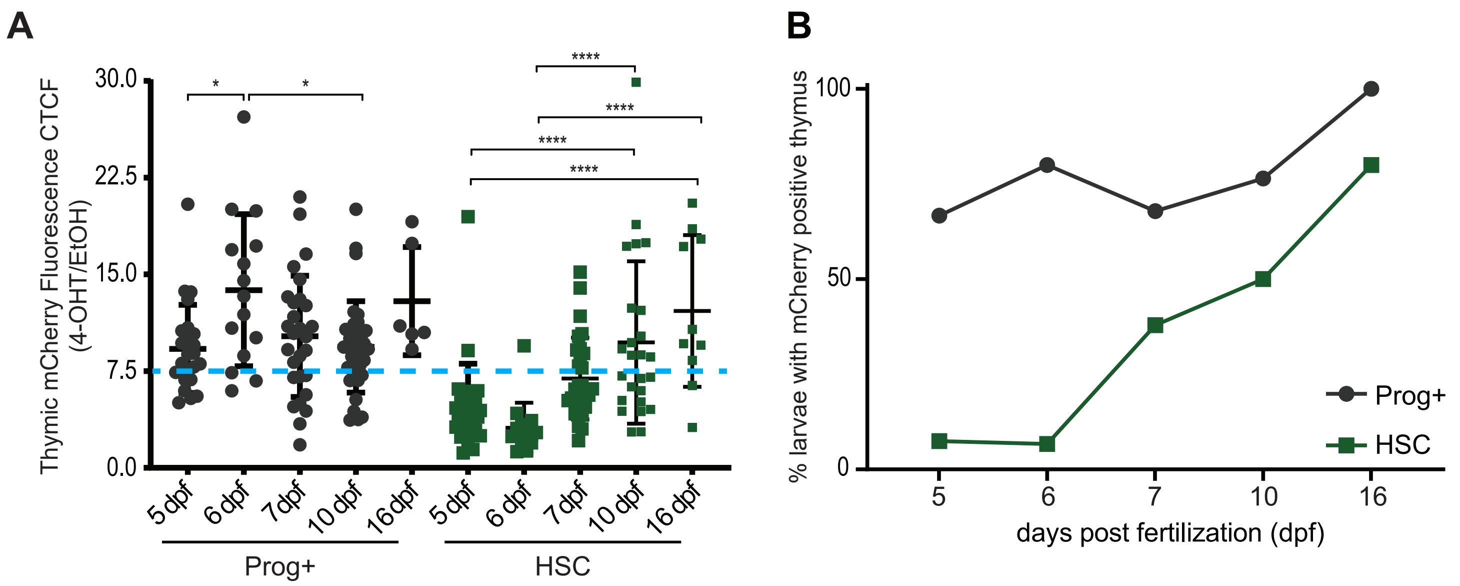


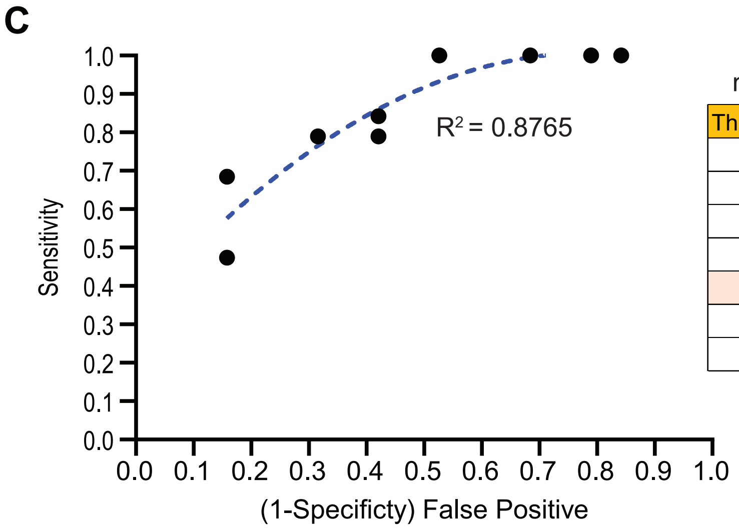


mCherry Fluorescence Threshold

| Threshold | Sensitivity | (1-Specificity) |
|---|---|---|
| 4.000 | 1.000 | 0.684 |
| 5.000 | 1.000 | 0.526 |
| 6.000 | 0.842 | 0.421 |
| 7.000 | 0.789 | 0.421 |
| 7.500 | 0.789 | 0.316 |
| 9.250 | 0.684 | 0.158 |
| 11.000 | 0.474 | 0.158 |

**Supplement Figure 3**

**Figure S3, related to Figure 3**. Supplemental lymphoid lineage tracing quantification for Prog+ and HSC populations. (A) Quantification of mCherry$^+$ thymic fluorescence from Prog+ and HSC lineage tracing cohorts. CTCF values for 4-OHT-treated embryos were normalized to ethanol controls. CTCF positive threshold of 7.5 (blue dotted line) was derived from a blinded analysis and demarcates the point at which mCherry$^+$ fluorescence is visibly detected in the thymus. Two-way ANOVA with Sidak's multiple comparison was used for analysis. N = 6-30 per larvae/day. Plots are individual data points for each biological replicate with mean ± standard deviation. *p-value $< 0.05$, ****p-value $\leq 0.0001$. (B) Quantification of the percentage of larvae with mCherry$^+$ thymus based on fluorescence imaging and the threshold defined in S3A. (C) Receiving Operating Curve (ROC) was used to calculate a fluorescence threshold (7.5) demarcating the thymic mCherry$^+$ fluorescence level for visual detection. Sensitivity and (1-Specificity) False Positive values were based on blinded scoring of 34 thymus images.

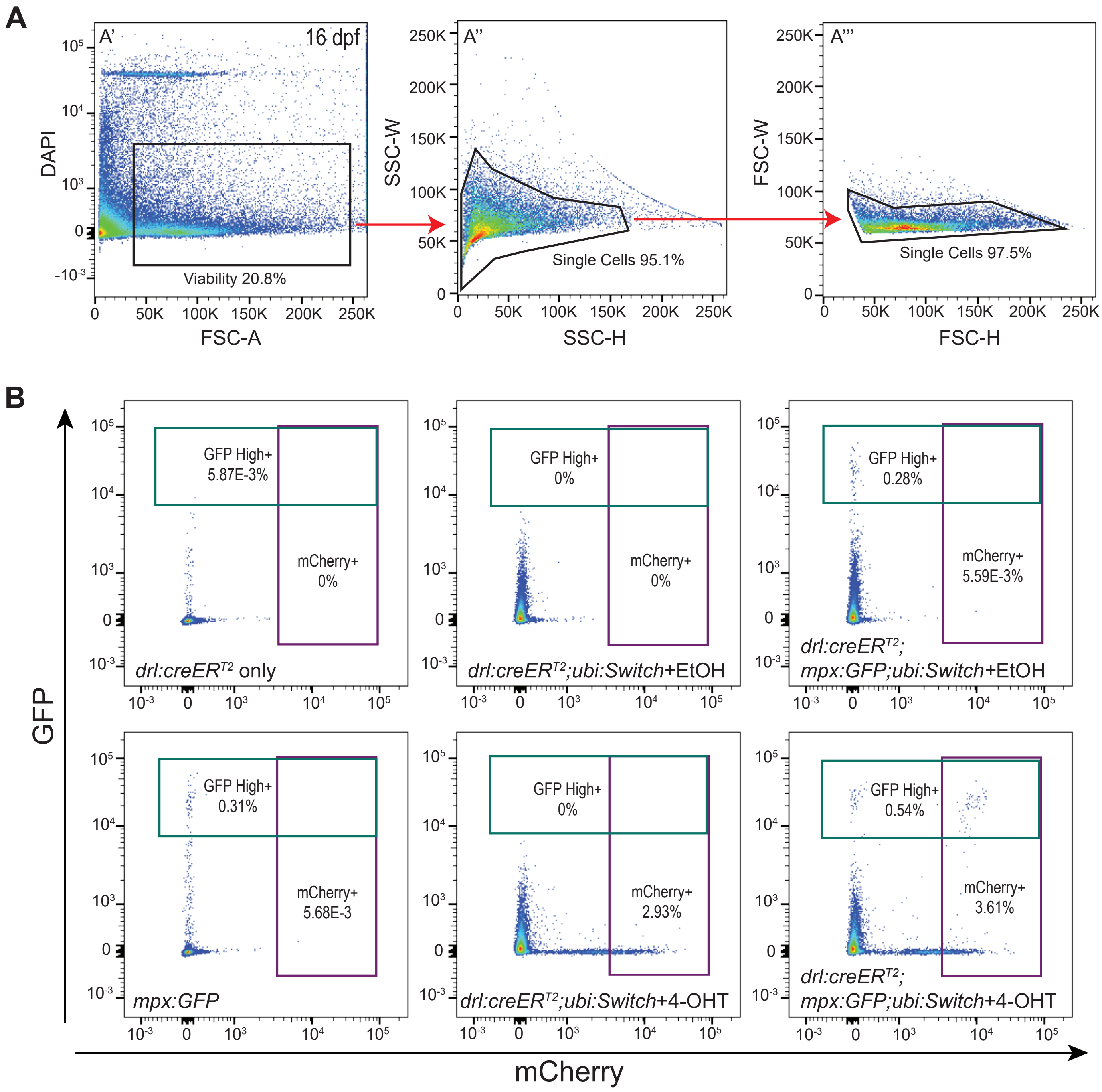


**Supplemental Figure 4**

**Figure S4, related to Figure 4**. Supplemental myeloid lineage tracing gating strategy for Prog+ and HSC populations. (A) Flow cytometry plots showing gating strategy for quantifying mCherry$^{+}$ Prog+ and HSC lineage-traced cells. Live cells were selected based on lack of DAPI signal (A'), then single cells were selected using SSC-W vs SSC-H (A'') and FSC-W vs FSC-H (A'''). (B) Flow cytometry plots showing gating strategy for the *mpx:GFP*$^{high}$ fraction and mCherry$^{+}$ switch cells based on control groups. *mpx:GFP*$^{high}$ negative controls: *Tg(drl:creER*$^{T2}$*), Tg(drl:creER*$^{T2}$*;ubi:Switch)* with 4-OHT or EtOH*;* switch negative controls: *Tg(drl:creER*$^{T2}$*; ubi:Switch)* and *Tg(drl:creER*$^{T2}$*;mpx:GFP;ubi:Switch)* with EtOH; *mpx:GFP*$^{high}$ positive controls: *Tg(mpx:GFP)* and *Tg(drl:creER*$^{T2}$*;mpx:GFP;ubi:Switch)* with EtOH; and switch only positive controls: *Tg(drl:creER*$^{T2}$*;ubi:Switch)* with 4-OHT.

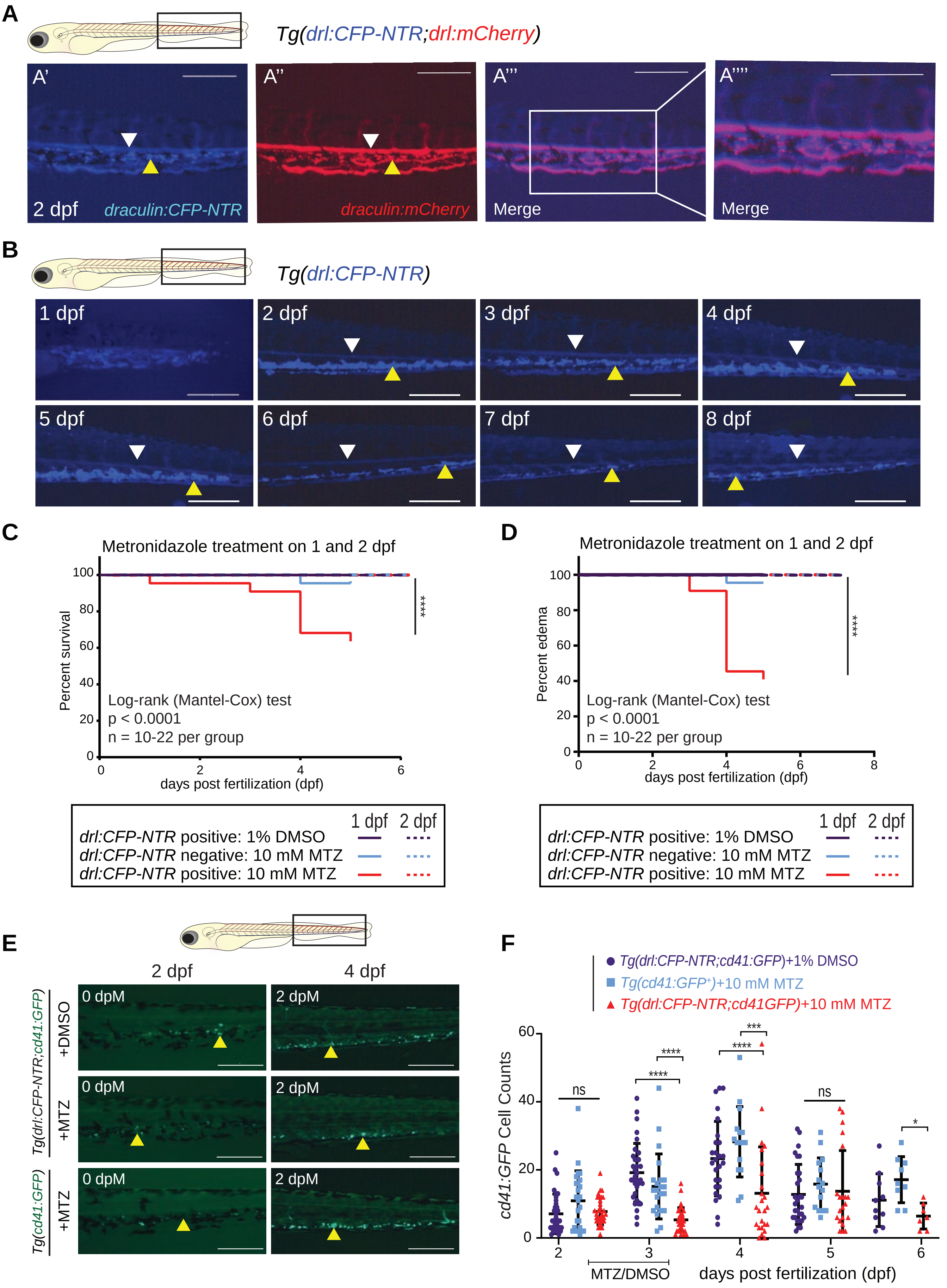


**Supplemental Figure 5.**

**Figure S5, related to Figure 5**. Supplemental information for the establishment of *in situ* HSC regeneration assay using NTR/MTZ system in zebrafish. (A) Fluorescent images showing the expression of *drl:CFP-NTR* (A’), *drl:mCherry* (A’’), and merged (A’’’) at 2 dpf. 1.75x inset in (A’’’’) indicating co-expression in both circulating (white arrowheads) and stationary (yellow arrowheads) cells. Scale bar = 500 μm. (B) Fluorescent images showing the expression pattern of *drl:CFP-NTR* within the CHT of 1–8 dpf zebrafish. Scale bar = 500 μm. (C) Kaplan-Meier curve for the survival of embryos treated with 1% DMSO or 10 mM MTZ for 20 hours at 1 dpf and 2 dpf. (D) Kaplan-Meier curve for the development of edema in embryos treated with 1% DMSO or 10 mM MTZ for 20 hours at 1 dpf and 2 dpf. Graphs show survival/edema development of *drl:CFP-NTR* positive + 1% DMSO (purple), *drl:CFP-NTR* negative + 10 mM MTZ (light blue), and *drl:CFP-NTR* positive + 10 mM MTZ (red). Log-rank (Mantel-Cox) test used for statistical analysis at 1 and 2 dpf (N = 10-22 per treatment group, ****p-value $\leq$ 0.0001). The solid lines are 1 dpf data and dotted lines are 2 dpf. (E) Fluorescent images of *Tg(drl:CFP-NTR;cd41:eGFP*$^+$*)* and *Tg(cd41:GFP*$^+$*)* embryos controls and experimental group at 2 and 4 dpf (0 and 2 dpM, respectively). Yellow arrowhead indicates stationary GFP$^+$ CHT cells. Scale bar = 500 μm. (F) Quantification of *cd41:GFP* fluorescence cell counts in *Tg(drl:CFP-NTR*$^+$*;cd41:GFP*$^+$*)* and *Tg(cd41:GFP+)* embryos treated with 10 mM MTZ or 1% DMSO (n = 3-15) from fluorescence images as shown in (E). Two-way ANOVA with Tukey’s multiple comparisons test was used for all statistical analyses. Plots are individual data points for each biological replicate with mean ± standard deviation. *p-value < 0.05, ***p-value $\leq$ 0.001, ****p-value $\leq$ 0.0001.
